# Venetoclax Resistance Creates Canonical Pyroptosis Vulnerability in AML

**DOI:** 10.64898/2026.10.14.759155

**Authors:** Qiuxia Wang, Shaohua Xu, Jizhen Cai, Qian Peng, Shuting Cao, Yanyan Zhang, Pengfei Lu, Xu Guo, Luzheng Liu, Wenna Fu, Song Shao, Jiazhuo Zhu, Bowen Qing, Hongli Li, Kang Liu, Kaibo Zhu, Hui Yang, Xinyu Yang, Ganqian Zhu

## Abstract

Venetoclax has transformed acute myeloid leukemia (AML) therapy, yet resistance is almost inevitable. Its diverse, often difficult-to-target drivers frequently converge on monocytic differentiation, a state that survives on MCL1 rather than BCL2. Here, we show that the very cells that escape BCL2 inhibition retain the canonical pyroptosis machinery of normal monocytes. Across 15 bulk and 4 single-cell cohorts, the canonical pyroptosis program was enriched in monocytic AML and monocytic leukemia stem cells, correlated with venetoclax resistance, and associated with inferior survival specifically after venetoclax-based therapy. In 24 cell lines and 142 patient specimens, both DPP8/9 inhibition by Val-boroPro and NLRP3 activation by lipopolysaccharide (LPS) plus nigericin preferentially killed monocytic AML cells through GSDMD-mediated pyroptosis, largely sparing non-monocytic AML and lymphoid leukemias. Venetoclax preferentially depleted primitive AML blasts but enriched a monocytic residuum vulnerable to canonical pyroptosis; combining venetoclax with Val-boroPro achieved deeper *ex vivo* clearance than either agent alone. In venetoclax-resistant xenografts, Val-boroPro suppressed leukemia progression, and adding venetoclax further reduced leukemia burden and prolonged survival beyond either monotherapy. Thus, venetoclax resistance itself exposes an alternative, druggable cell-death program that can be therapeutically exploited to overcome this resistance.

**Key Points:**

1. Monocytic AML retains canonical pyroptosis program, creating a targetable vulnerability linked to venetoclax resistance.
2. Venetoclax enriches Val-boroPro-sensitive monocytic residual cells; their combination achieves deeper clearance and prolongs survival.

## Introduction

By inhibiting BCL2, a critical survival dependency of acute myeloid leukemia (AML) blasts and leukemia stem cells (LSCs),^1,2^ venetoclax (VEN) became the first approved AML agent to directly target apoptosis when combined with hypomethylating agents (HMAs)^3,4^ or low-dose cytarabine,^5,6^ transforming treatment for the ∼50% of newly diagnosed patients considered unfit for intensive chemotherapy,^7^ who have long faced a dismal prognosis.^8^ VEN is also now being incorporated into intensive induction for fit patients.^9–12^ However, ∼33% of patients receiving VEN-based lower-intensity therapy fail to achieve complete remission,^4^ most responders relapse,^13^ and options after VEN plus HMA failure remain scarce.^14^ Thus, primary and acquired resistance are principal barriers to durable benefit.

VEN resistance is multifactorial: (i) altered mitochondrial apoptotic priming driven by upregulation of alternative anti-apoptotic BCL2 family proteins such as MCL1,^15–19^ BCL-xL,^15,18–21^ and BFL1;^15,22^ (ii) recurrent *TP53*,^20,23^ *NRAS/KRAS*,^24^ *PTPN11*,^15^ *FLT3*^25^ mutations or acquired *BAX* mutations,^26^ several embedded in the ELN2024 risk stratification for VEN plus HMA regimen;^25^ (iii) disrupted mitochondrial homeostasis.^27–30^ Although combining FLT3 inhibition with VEN has shown clinical activity,^14,31^ efforts to cotarget other resistance drivers such as MCL1, BCL-xL, or mutant TP53 are likely to be ultimately constrained by cardiotoxicity, thrombocytopenia, and severe adverse events, respectively.^14^

Strikingly, diverse resistance mechanisms tend to converge on a single phenotype: monocytic differentiation. Both FAB M4/M5 disease^15,19,32–36^ and VEN-driven clonal selection of monocytic LSCs that regenerate monocytic AML^33,37^ are established non-driver resistance phenotypes, and the very *NRAS*, *KRAS*, *PTPN11* and *FLT3* mutations that drive resistance are enriched in monocytic disease.^38–40^ Their unifying basis is a metabolic and survival dependency on MCL1 rather than BCL2, shared by monocytic blasts, monocytic LSCs and normal monocytes,^14,15,33,34,37^ rendering these cells intrinsically refractory to BCL2 inhibition. Moreover, given the aggressive tempo of AML, selecting pre-existing clones harboring resistance drivers that predispose them to monocytic differentiation is a more efficient evolutionary route to resistance than acquiring *de novo* mutations.^14,33^ We therefore reasoned that a superior strategy would not contest upstream drivers but instead eliminate monocytic AML cells by exploiting a lineage property retained after leukemic transformation.

Yet apoptosis remains the only programmed cell death directly harnessed by approved cancer therapies, leaving pyroptosis clinically unexploited. Mature myeloid cells, including monocytes, macrophages, granulocytes, and dendritic cells (DCs), are equipped with pattern recognition receptors (PRRs) that, upon sensing microbe-or danger-associated molecular patterns (MAMPs/DAMPs), assemble canonical inflammasomes, triggering proximity-induced autoproteolysis of caspase-1 and cleavage of gasdermin D (GSDMD) to execute canonical pyroptosis, a lytic pro-inflammatory cell death.^41–43^ Unlike the GSDME- and GSDMB-mediated pyroptotic programs that tumors routinely silence,^44–46^ canonical GSDMD pyroptosis acts as an antimicrobial effector rather than a tumor-suppressive barrier^44,46^ and may therefore be preserved through malignant transformation, offering a driver-independent vulnerability. Consistent with this, the DPP8/9 inhibitor Val-boroPro (VBP), the first small molecule identified to induce canonical pyroptosis in monocytes and macrophages,^47^ was subsequently shown to trigger CARD8 inflammasome-mediated pyroptosis in a subset of AML cell lines,^48^ prompting phase 1 clinical evaluation (NCT05703542) in relapsed/refractory (R/R) AML and MDS without biomarker-based patient selection.^49^ However, because AML cells are arrested at varied stages of myeloid differentiation whereas canonical pyroptosis is a property of mature myeloid cells,^8,41–43^ it remains unknown which AML subtypes are poised for pyroptotic execution, whether this susceptibility is coupled to VEN resistance, and whether engaging it can overcome VEN resistance by eradicating monocytic cells.

Here, we show that monocytic AML retains canonical pyroptosis machinery at levels comparable to normal monocytes. Expression of this program is associated with VEN resistance and inferior outcomes after VEN-based but not conventional therapy. *Ex vivo* profiling of 106 primary AML samples revealed reciprocal sensitivity to VEN and canonical pyroptosis induction: VBP and the NLRP3 stimulus plus nigericin (LPS+Nig) preferentially killed monocytic and VEN-resistant AML cells. Critically, VEN preferentially depleted non-monocytic blasts while enriching a VBP-sensitive monocytic residuum, creating a dynamic vulnerability exploited by the VBP+VEN combination to achieve superior efficacy in VEN-resistant xenografts. These findings identify canonical pyroptosis as a targetable vulnerability of monocytic AML and support incorporating pyroptosis-inducing agents into VEN-based regimens.

## Methods

### Cell culture

AML and acute lymphoblastic leukemia (ALL) cell lines were cultured in RPMI 1640 with 10% fetal bovine serum. VEN-resistant lines were generated by stepwise VEN escalation.

### Primary leukemia specimens

Bone marrow (BM) specimens from 106 patients with AML and 36 with ALL were obtained at Xiangya Hospital with written informed consent under ethics approval and in accordance with the Declaration of Helsinki.

### Drug treatments and assays

Cell-line dose-response assays used VBP or VEN for 48 h. Primary specimens received VEN (0.5 μM, 36 h), VBP (5 μM, 24 h), or VEN+VBP, with VBP added during the final 24 h. LPS+Nig comprised LPS (10 μg/mL, 24 h) followed by nigericin (10 μM, 1 h). Assays included CCK-8, flow cytometry, LDH release, and immunoblotting.

### Xenograft studies

After 100 cGy total-body irradiation, NCG mice received VEN-resistant MV4-11 cells or primary FAB M5 AML BM mononuclear cells intravenously. Mice were randomized to vehicle, VEN, VBP, or VEN+VBP. VEN was administered by oral gavage and VBP intraperitoneally 5 days/week for 4 weeks. Leukemia burden was assessed by hCD45⁺ cells in peripheral blood and BM. Animal studies were approved by Hunan University’s Animal Care and Welfare Committee.

### Transcriptomics

Bulk and single-cell analyses are detailed in supplemental Methods; cell-line RNA-seq reads are publicly available in SRA (PRJNA1516960).

### Use of artificial intelligence tools

ChatGPT (OpenAI) and Claude (Anthropic) were used solely to improve grammar and language in author-generated text; authors reviewed all revisions.

**Additional methods are provided in the supplemental Methods.**

## Results

### 1. Monocytic differentiation confers canonical pyroptotic vulnerability in AML

A previous study showed that VBP triggers canonical pyroptotic cell death (inflammasome-dependent, GSDMD-mediated) in a subset of AML cell lines.^48^ Indeed, in pan-cancer analyses of the TCGA and CCLE datasets, AML showed the highest expression of canonical pyroptosis machinery, including PRRs, caspases, and pore-forming GSDMD,^41–43^ albeit with a notable proportion of low-expressing cases (Figure 1A; supplemental Figure 1A). Given that canonical pyroptosis is an innate feature of mature myeloid cells, we hypothesized that pyroptotic propensity is linked to AML differentiation state.

**Figure 1.**
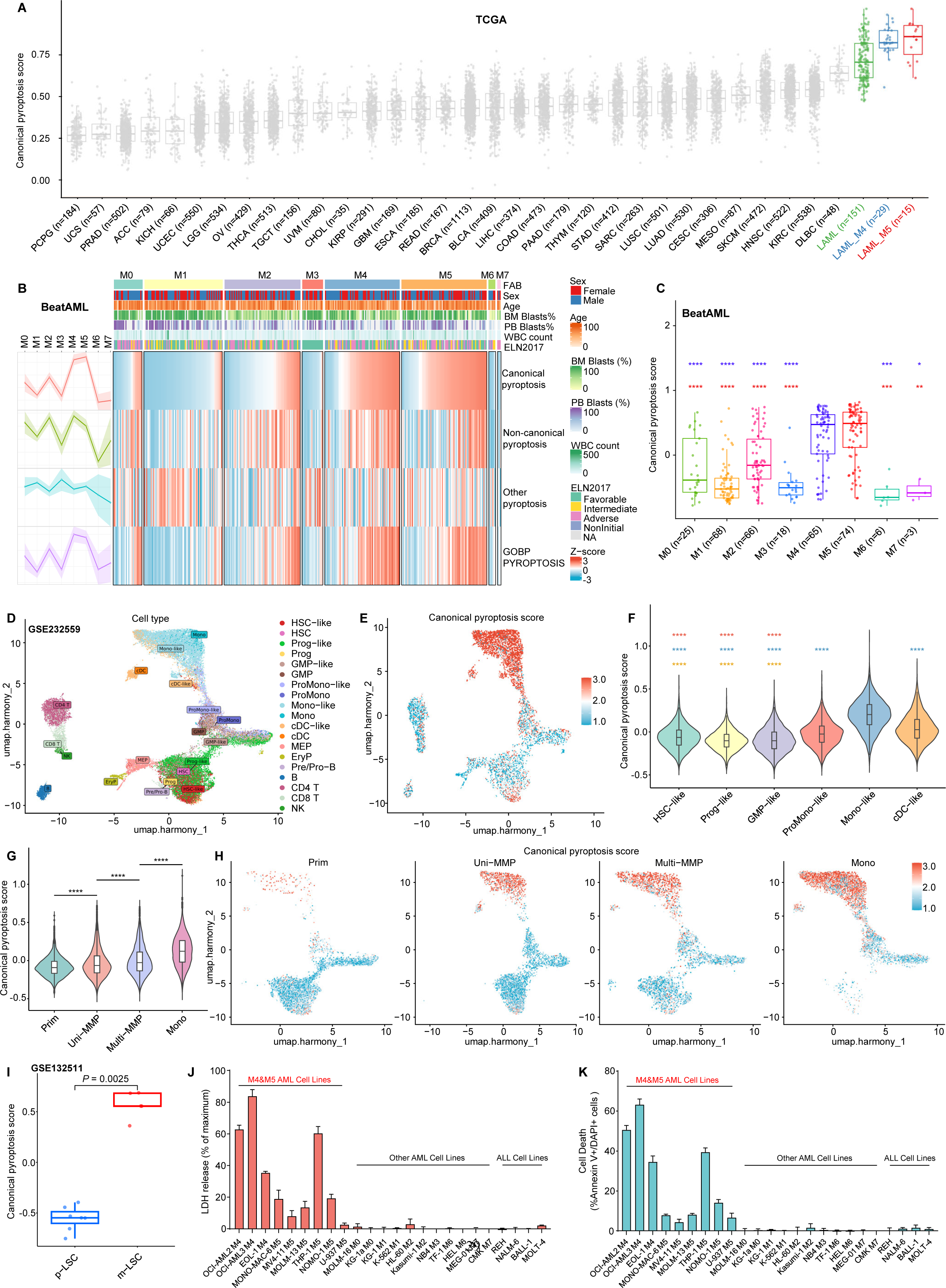
Monocytic differentiation confers canonical pyroptotic vulnerability in AML. **(A)** Box plots showing canonical pyroptosis signature scores across TCGA tumor types, with acute myeloid leukemia (AML) and its FAB M4 and M5 subsets highlighted. **(B)** Heat map depicting the pyroptosis-related gene signature scores across FAB subtypes in the BeatAML cohort. Pyroptosis-associated genes were grouped into canonical pyroptosis, non-canonical pyroptosis and other-pyroptosis signatures, together with the GOBP_PYROPTOSIS gene set. Samples are ordered according to FAB classification, with annotations for sex, age, bone marrow (BM) blast percentage, peripheral blood (PB) blast percentage, white blood cell (WBC) count and European LeukemiaNet 2017 (ELN2017) risk stratification. Marginal line plots on the left summarize trends in the mean score of each pyroptosis signature across FAB subtypes. Colors represent row-wise Z-score-normalized signature/gene-set scores. **(C)** Canonical pyroptosis signature scores across FAB subtypes in the BeatAML cohort. Violet-blue and red asterisks denote comparisons with M4 and M5, respectively. **(D)** Uniform manifold approximation and projection (UMAP) visualization of 32,319 cells from 27 samples obtained from 25 AML patients in the GSE232559 scRNA-seq dataset, colored by annotated cell types identified based on canonical marker gene expression. **(E)** UMAP visualization of canonical pyroptosis signature scores for the cells shown in (D). **(F)** Violin plots comparing canonical pyroptosis signature scores across AML cell subpopulations in 24,996 AML cells from 27 samples in the GSE232559 scRNA-seq dataset. Colored asterisks above individual populations indicate selected pairwise comparisons with the color-matched reference populations: red, ProMono-like; blue, Mono-like; and orange, cDC-like. **(G)** Violin plots showing canonical pyroptosis signature scores in 23,317 AML cells from 25 samples in the GSE232559 scRNA-seq dataset, grouped by differentiation state: predominantly primitive AML (Prim), predominantly monocytic AML (Mono), and two mixed monocytic-primitive subtypes defined by leukemia stem cell (LSC) activity distribution: Uni-MMP, with LSC activity restricted to primitive cells, and Multi-MMP, with LSC activity present in both primitive and monocytic cells. Groups are ordered by increasing monocytic differentiation. **(H)** UMAP visualization of canonical pyroptosis signature scores in the same 23,317 AML cells as in (G), stratified by differentiation-state group. **(I)** Canonical pyroptosis signature scores in primitive LSCs (p-LSCs) versus monocytic LSCs (m-LSCs) from the GSE132511 dataset. **(J-K)** LDH release (J) and flow-cytometric quantification of Annexin V⁺DAPI⁺ cell death (K) in AML and ALL cell lines following canonical pyroptosis induction by LPS priming (10 μg/mL, 24 h) followed by nigericin treatment (10 μM, 1 h). In (A), each dot represents one TCGA tumor sample; in (C) and (I), each dot represents one patient sample. Statistical significance in (C, F, G) and (I) was determined by two-sided Wilcoxon rank-sum tests for indicated comparisons. \**P* < .05, \*\**P* < .01, \*\*\**P* < .001, \*\*\*\**P* < .0001. Data in (J) and (K) are shown as mean ± s.d. from three independent experiments.

To identify AML subtypes primed for pyroptosis, we profiled distinct pyroptosis programs across differentiation states defined by the FAB classification,^32^ which lacks independent prognostic value in conventional chemotherapy after genetic alterations are accounted for.^50^ In BeatAML,^51^ myelomonocytic/monocytic (M4/M5) AML showed pronounced enrichment of canonical pyroptosis gene expression (Figure 1B), with significantly higher signature scores than other FAB subtypes, including primitive (M0/M1), granulocytic (M2/M3), erythroid (M6), and megakaryoblastic (M7) (Figure 1C). By contrast, genes associated with non-canonical pyroptosis (inflammasome-independent, GSDMD-mediated) or other-pyroptosis (other GSDMs-mediated) showed no subtype-specific trend (Figure 1B). Similar results were observed in the pediatric and young-adult TARGET-AML cohort (supplemental Figure 1B-C). Beyond these 2 largest AML datasets, after stringent batch-effect correction (supplemental Figure 2A, C, E), integration of 12 additional large-scale RNA-seq, GPL96 microarray, and GPL570 microarray datasets yielded consistent results (supplemental Figure 2B, D, F).

Given the pronounced inter- and intra-patient heterogeneity and the complex tumor microenvironment of AML, to resolve canonical pyroptosis activity at cellular resolution, we analyzed a single-cell RNA-seq dataset (GSE232559) comprising 27 AML samples classified by blast differentiation stage as Prim-, Uni-MMP-/Multi-MMP-, and Mono-AML,^37^ and further integrated three independent single-cell RNA-seq datasets (GSE185381, GSE235063, and GSE239721; supplemental Figure 3A). UMAP displayed cells across patients with annotated types defined by marker expression (Figure 1D; supplemental Figure 3B-D). Canonical pyroptosis activity was heterogeneous across cell populations (Figure 1E; supplemental Figure 3E). Within AML cells, differentiated ProMono-like, Mono-like, and cDC-like populations showed significantly higher activity than primitive HSC-like, Prog-like, and GMP-like populations (Figure 1F; supplemental Figure 3F). Notably, although malignant and incompletely differentiated, monocytic-like AML cells expressed canonical pyroptosis genes at levels comparable to normal monocytes, suggesting preserved pyroptotic capacity (supplemental Figure 3F). Consistently, pooled malignant cells showed a stepwise increase of canonical pyroptosis activity across Prim-, Uni-MMP-, Multi-MMP-, and Mono-AML, peaking in Mono-AML and corroborating the bulk RNA-seq and microarray results (Figure 1G-H). Given the heterogeneity and repopulating capacity of LSCs, we also compared primitive LSCs (p-LSCs, which initiate primitive AML) with monocytic LSCs (m-LSCs, which initiate monocytic AML).^33,37^ Bulk RNA-seq of sorted LSCs (GSE132511) showed significantly higher canonical pyroptosis pathway activity in m-LSCs than in p-LSCs (Figure 1I).

Together, these results indicate that AML cells with monocytic differentiation, like normal monocytes, constitutively express canonical pyroptosis genes and may therefore be more readily induced to undergo pyroptosis than immature, non-monocytic, or non-myeloid cells. Such cells predominate in M4/M5 AML but also occur in a minority of other FAB subtypes.

To confirm this differentiation- and lineage-specific vulnerability, we induced pyroptosis in 20 AML cell lines grouped by FAB subtype and 4 ALL cell lines. LPS+Nig, a commonly used stimulus for NLRP3-dependent pyroptosis *in vitro*, induced robust LDH release and cell death only in M4/M5 AML lines; immature, granulocytic, erythroid, megakaryocytic, and lymphoid lines were largely non-responsive (Figure 1J-K). Most nonviable cells after LPS+Nig treatment were Annexin V/DAPI double-positive (supplemental Figure 4A), consistent with pyroptotic membrane perforation.

Collectively, these findings indicate that monocytic AML expresses sufficient canonical pyroptosis machinery to prime them for pyroptosis induction.

### 2. Monocytic AML is preferentially susceptible to Val-boroPro-induced pyroptosis

As a nonselective small-molecule inhibitor of S9 post-proline-cleaving serine proteases (DPP4, DPP7, DPP8, DPP9 and FAP), VBP has been shown to trigger AML pyroptosis through DPP8/9 inhibition-induced CARD8 inflammasome assembly and subsequent GSDMD cleavage.^47,48^ Unlike LPS+Nig, this single agent induces a truncated canonical pyroptosis program without substantial NF-κB-primed transcription of inflammatory factors, potentially making it more practical for AML treatment. Indeed, VBP is already in a clinical trial for R/R AML and MDS,^49^ but the determinants of response remain unclear. We therefore evaluated the aforementioned cell lines panel. Like LPS+Nig, VBP inhibited M4/M5 cell-line growth with nanomolar potency (Figure 2A), whereas all tested M0-M3, M6, M7, and B/T lymphoid lines were highly resistant even at 100 μM (Figure 2B). VBP-induced AML cell death likewise manifested as dose-dependent Annexin V^+^/DAPI^+^ staining and LDH release, predominantly in M4/M5 lines (Figure 2C-D; supplemental Figure 4B). Cleavage of GSDMD into its active N-terminal fragment (GSDMD-N) was detected in the M4/M5 lines responsive to VBP or LPS+Nig (Figure 2E).

**Figure 2.**
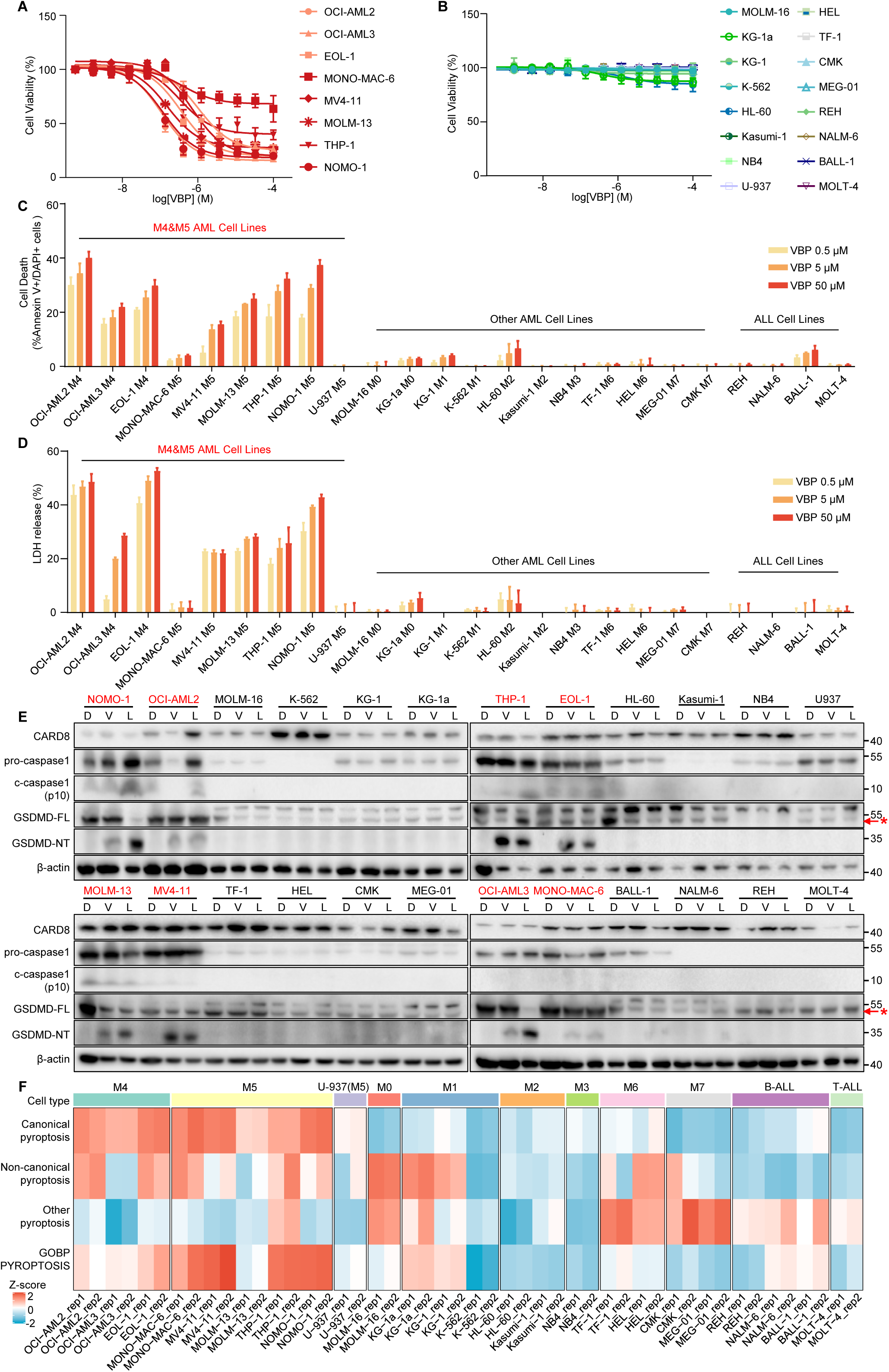
VBP selectively induces canonical pyroptotic cell death in monocytic AML cells. **(A)** Dose-response curves of cell viability in VBP-sensitive FAB M4/M5 AML cell lines treated with VBP for 48 h. **(B)** Dose-response curves of cell viability in VBP-resistant AML and ALL cell lines treated with VBP for 48 h. VBP-sensitive and -resistant cell lines were classified according to their cell-viability dose-response profiles across the tested VBP concentrations. **(C)** Flow-cytometric quantification of VBP-induced cell death in AML and ALL cell lines treated with increasing concentrations of VBP for 48 h. **(D)** Percentage of maximal LDH release from AML and ALL cell lines treated with increasing concentrations of VBP for 48 h. **(E)** Immunoblot analysis of CARD8, pro-caspase-1, cleaved caspase-1, full-length GSDMD (GSDMD-FL), and the cleaved GSDMD N-terminal fragment (GSDMD-N) in the indicated AML and ALL cell lines following treatment with DMSO (D), VBP (V), or LPS plus nigericin (L). β-actin was used as a loading control. The red asterisk and red arrow mark the specific band of GSDMD-FL. Red labels denote the VBP-sensitive M4/M5 AML cell lines shown in (A). **(F)** Heat map depicting the pyroptosis-related gene signature scores across AML and ALL cell lines. Genes were grouped into canonical pyroptosis, non-canonical pyroptosis and other-pyroptosis signatures, together with the GOBP_PYROPTOSIS gene set. Cell lines are ordered by FAB subtype or lymphoid lineage, with U-937 annotated separately. Colors represent row-wise Z-score-normalized signature/gene-set scores. In (A) and (B), cell viability was normalized to matched DMSO-treated controls. In (C) and (D), values are shown after subtraction of the matched DMSO-control baseline. Data in (A-D) are shown as mean ± s.d. from three independent experiments.

HL-60 is an atypical M2 AML cell line known to spontaneously differentiate beyond the myeloblast stage into promyelocytes, myelocytes, and mature granulocytes,^52^ potentially explaining its minor but discernible VBP-induced pyroptotic cell death (Figure 2C). Because only a small fraction differentiates and the fate of these cells is either death or exiting cell cycle, the bulk culture remained resistant to VBP (Figure 2B). Consistent with previous work,^47^ cleaved caspase-1 was not captured in most responsive cells after VBP or LPS+Nig treatment, except LPS+Nig-treated NOMO-1, OCI-AML2, and THP-1 cells (Figure 2E), likely because the gradual release of cellular contents allows thorough extracellular degradation.^53^ Actually, LPS+Nig elicits weaker pyroptosis in monocytes than in terminally differentiated macrophages (e.g., PMA-differentiated THP-1 cells), reflecting the lower expression of pyroptosis machinery in monocytes. Moreover, VBP-induced pyroptosis is slower than that induced by LPS+Nig in both monocytes and macrophages (24-48 h vs. 0.5-1 h).^47,54,55^ Accordingly, after VBP treatment, a weak cleaved caspase-1 band was detectable only in GSDMD-deficient THP-1 monocytes, in which impaired membrane perforation permitted its intracellular accumulation.^53^ Nevertheless, although immunology studies favor vigorous pyroptosis with robust inflammatory factor release, the VBP induced pyroptosis is sufficient to reduce monocytic AML growth (Figure 2A).

Although CARD8, caspase-1, and GSDMD are core components of VBP-induced pyroptosis, their expression only partially determined responses to VBP and LPS+Nig (Figure 2E). To define the molecular basis of differential susceptibility and its relationship to myeloid differentiation, we performed RNA-seq across all cell lines. Consistent with the protein data, *CARD8* was expressed in most resistant lines (supplemental Figure 5A). In contrast, sensitive M4/M5 lines expressed canonical pyroptosis genes at higher levels than resistant lines (Figure 2F; supplemental Figure 5A-B), had significantly higher monocytic and granulocytic differentiation scores, and had lower stemness scores (supplemental Figure 5C-E). Notably, genes encoding DAMP-sensing receptors such as P2X7, NLRP3 and AIM2^44^ were also highly expressed in monocytic AML cells (supplemental Figure 5A), potentially allowing DAMPs released by initial VBP-induced pyroptosis to engage these receptors and amplify pyroptosis. Thus, responses to VBP or LPS+Nig reflect broad expression of the canonical pyroptosis gene set rather than a few individual genes such as *CARD8* or *CASP1*.^48^

Unlike other monocytic AML lines, U-937 was established from diffuse histiocytic lymphoma^52^, an ambiguous entity subsequently shown in most cases to be lymphoid rather than myeloid.^56^ Beyond this unusual tissue origin, it also carries a *CALM*-*AF10* fusion, a rare but recurrent cytogenetic abnormality in T-ALL, AML, and acute leukemia of ambiguous lineage, suggesting lineage infidelity and potential transdifferentiation. Although its monocytic and granulocytic differentiation scores resembled those of other monocytic lines (supplemental Figure 5C-D), U-937 expressed fewer canonical pyroptosis genes (Figure 2F; supplemental Figure 5A-B), explaining its resistance to both LPS+Nig and VBP (Figure 1J-K; Figure 2B-E), which is also reported previously.^47,48^ U-937 is therefore a rare exception in which monocytic differentiation and canonical pyroptosis gene expression are uncoupled, reinforcing that gene expression directly determines pyroptotic vulnerability. Moreover, despite a conference abstract associating high *EPCAM* expression with VBP resistance,^57^ *EPCAM* expression was low in most non-responsive lines (supplemental Figure 5A, F).

Collectively, these data show that VBP, or other potential canonical pyroptosis inducers preferentially kill monocytic AML cells. The integrated gene signature, or the clinically more practical differentiation status, may predict this response.

### 3. Primary monocytic AML is intrinsically susceptible to canonical pyroptosis induction *ex vivo*

No large-scale data on AML sensitivity to VBP have been reported. To test whether monocytic differentiation sensitizes primary AML cells to canonical pyroptosis, we performed *ex vivo* drug-sensitivity profiling of bone marrow mononuclear cells (BMMNCs) from 106 newly diagnosed patients with AML spanning FAB subtypes (M0-M3, *n* = 53; M4, *n* = 12; M5, *n* = 41) and 36 patients with ALL. Samples with sufficient cellularity underwent parallel CCK-8 viability, Annexin V/DAPI, and LDH-release assays after VBP or LPS+Nig treatment (*n* = 107); samples with limited cell numbers underwent CCK-8 alone (*n* = 35).

An integrated heatmap showed enrichment of responsive samples in FAB M4/M5 AML (Figure 3A). After VBP treatment, CCK-8 and Annexin V/DAPI assays showed significantly lower viability in M4/M5 than in M0-M3 AML or ALL samples (Figure 3B-C), and VBP induced membrane rupture and LDH release predominantly in M4/M5 samples (Figure 3D; supplemental Figure 6). These response patterns were highly reproducible with LPS+Nig (Figure 3E-G; supplemental Figure 6). Unlike the all-or-none cell-line responses (Figure 1J-K; Figure 2A-E), a small fraction of non-monocytic M0-M3 samples was as sensitive to VBP or LPS+Nig as M4/M5 samples, consistent with high canonical pyroptosis gene expression in a minority of M0-M3 samples (Figure 1B-C; supplemental Figure 1B-C; supplemental Figure 2B, D, F). This reflects the heterogeneity and phenotypic plasticity of primary AML, although *ex vivo* culture-induced differentiation cannot be excluded.

**Figure 3.**
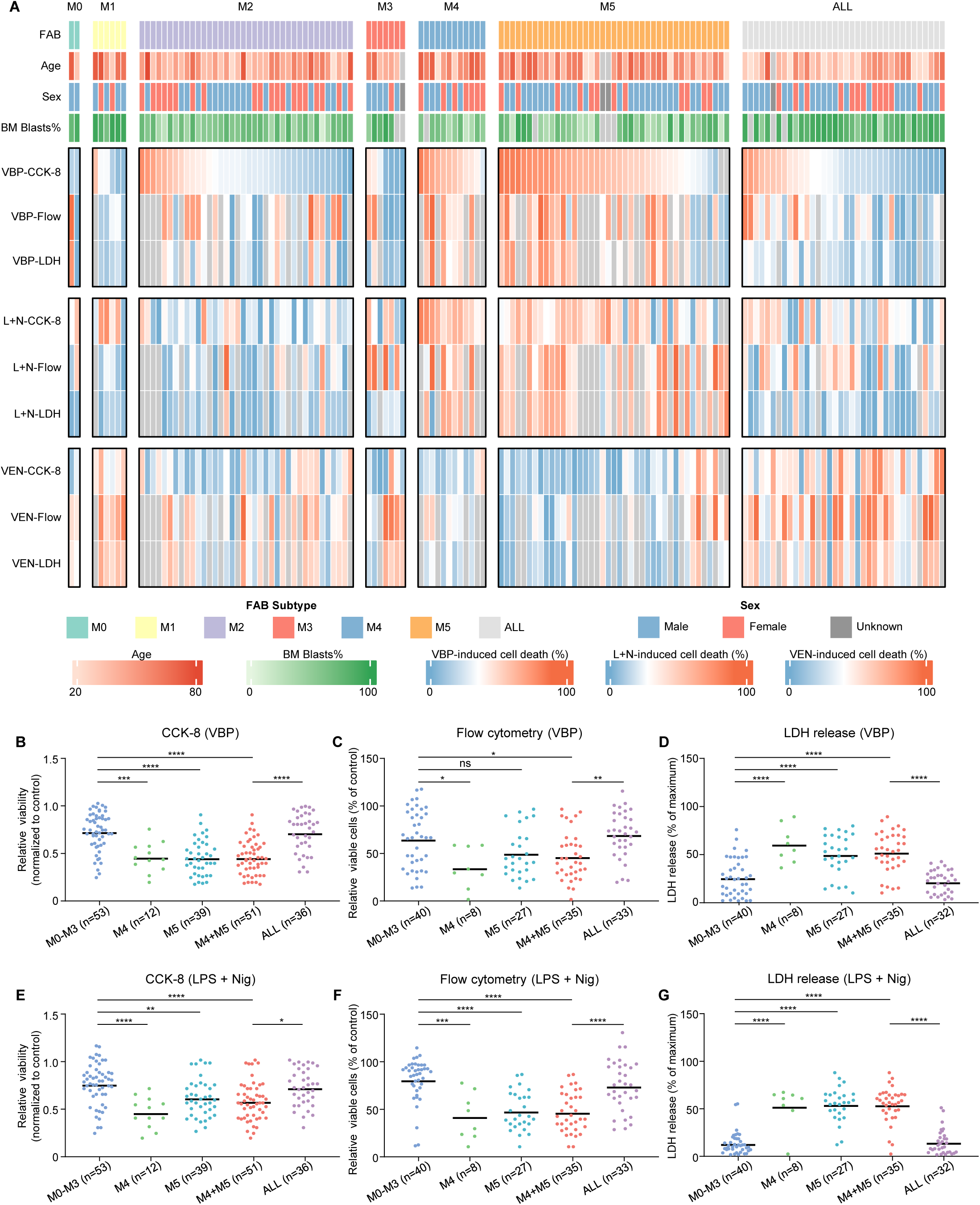
Primary monocytic AML specimens are selectively susceptible to canonical pyroptosis induction *ex vivo*. **(A)** Integrated heat map of *ex vivo* drug responses of bone marrow mononuclear cells (BMMNCs) from primary AML and ALL specimens. Samples include 106 AML specimens spanning FAB M0-M5 and 36 ALL specimens, ordered by FAB subtype or lymphoid lineage. Top annotations indicate disease classification, age, sex and BM blast percentage. The heat map shows responses to VBP, LPS plus nigericin (L+N) and VEN, assessed using CCK-8 viability, Annexin V/DAPI flow cytometry, and LDH release assays. Color gradients represent age, BM blast percentage or treatment-induced cell death, as indicated in the scale bars; grey denotes unavailable measurements. For visualization in (A), CCK-8- and flow-cytometry-derived relative viability values were converted to cell death as 100% minus relative viability, whereas LDH release values were corrected by subtraction of the matched vehicle-control baseline. **(B-D)** *Ex vivo* responses of primary AML and ALL specimens to VBP treatment, assessed by CCK-8 relative viability (B), Annexin V/DAPI flow cytometry relative viability (C) and baseline-corrected LDH release (D). **(E-G)** *Ex vivo* responses of primary AML and ALL specimens to LPS+Nig-induced canonical pyroptosis, assessed by CCK-8 relative viability (E), Annexin V/DAPI flow cytometry relative viability (F) and baseline-corrected LDH release (G). For *ex vivo* treatments, VEN was used at 0.5 μM for 36 h, VBP at 5 μM for 24 h, and LPS+Nig consisted of LPS priming at 10 μg/mL for 24 h followed by nigericin treatment at 10 μM for 1 h. In (B-G), each dot represents one primary specimen and horizontal lines indicate the mean. For (B, C, E) and (F), viability was normalized to matched DMSO-treated controls. For (D) and (G), LDH release was calculated as the percentage of maximal release and corrected by subtraction of the matched DMSO-control baseline. Assay-specific sample numbers varied according to available cellularity. Statistical significance was determined by one-way ANOVA followed by Tukey’s multiple-comparisons test for indicated comparisons; ns, not significant; \**P* < .05, \*\**P* < .01, \*\*\**P* < .001, \*\*\*\**P* < .0001.

Collectively, these findings confirm that primary monocytic AML cells not only express higher levels of canonical pyroptosis genes but also are highly susceptible to canonical pyroptosis induction.

### 4. Venetoclax resistance is associated with enhanced canonical pyroptosis susceptibility in AML

Multiple VEN-resistance drivers, such as *NRAS*, *KRAS*, *PTPN11* and *FLT3* mutations, confer a monocytic phenotype that is itself associated with VEN resistance.^14,24,38–40^ Given the association between pyroptosis gene expression and monocytic differentiation, we asked whether canonical pyroptosis susceptibility is also associated with VEN resistance. In BeatAML, a higher VEN area under the dose-response curve (AUC), indicating resistance, correlated positively with canonical pyroptosis gene expression (Figure 4A; supplemental Figure 7A), regardless of FAB subtype (supplemental Figure 7B-E). Consistently, VEN sensitivity, measured by selective drug sensitivity score (sDSS), correlated negatively with canonical pyroptosis activity in FIMM-AML (Figure 4B; supplemental Figure 8). As expected, the GOBP_PYROPTOSIS gene set score correlated with the canonical pyroptosis score and with VEN AUC/sDSS because this pan-pyroptosis set includes canonical pyroptosis genes (supplemental Figure 7, 8). By contrast, other-pyroptosis and non-canonical pyroptosis gene expression showed little association with VEN AUC/sDSS (supplemental Figure 7, 8). Importantly, elevated canonical pyroptosis gene expression was associated with inferior outcomes after VEN-based therapies (Figure 4C), but not in chemotherapy-treated BeatAML or TARGET-AML cohorts (supplemental Figure 9). Thus, high canonical pyroptosis gene expression is associated with poor efficacy of VEN-based therapy.

**Figure 4.**
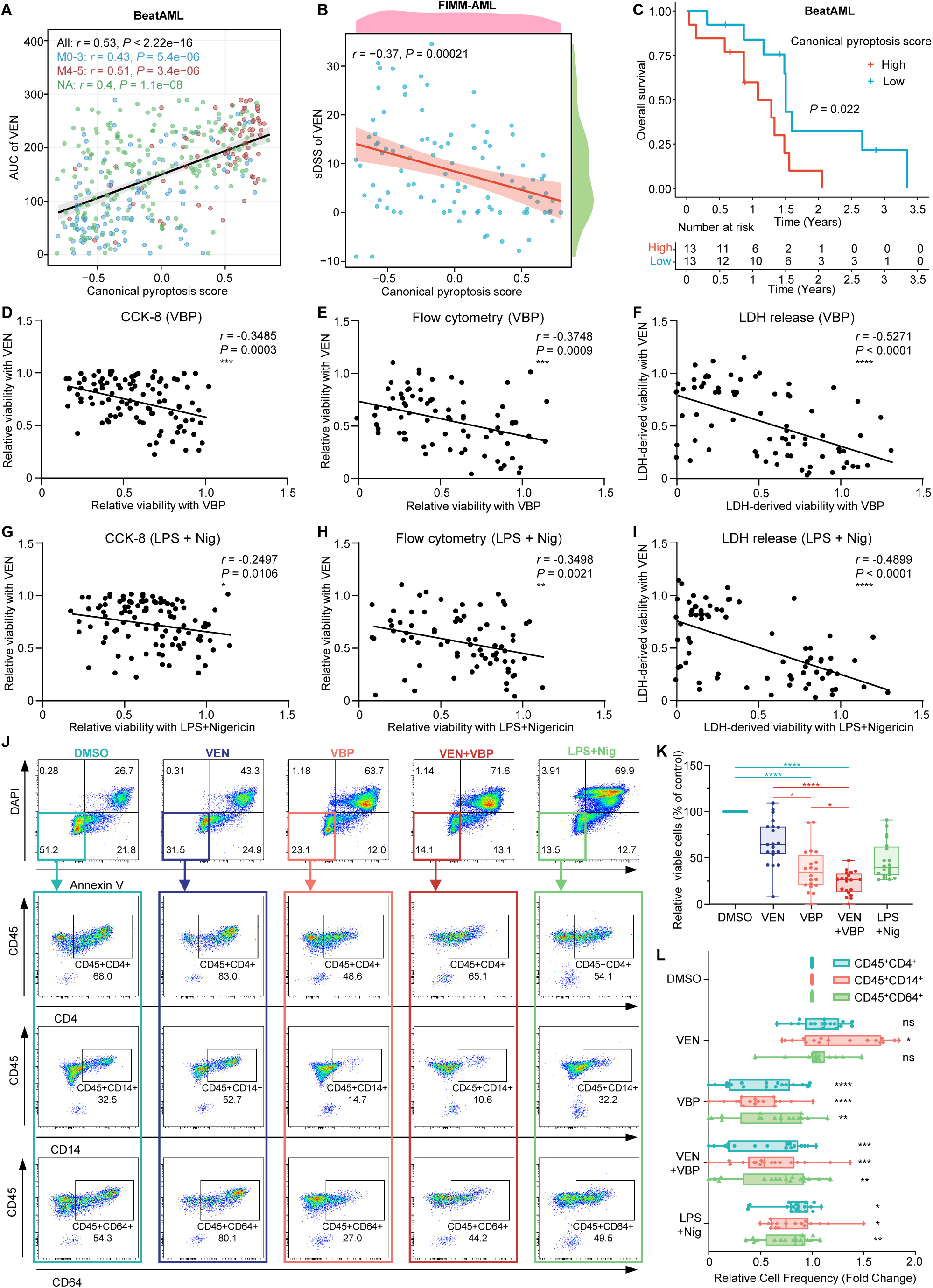
VEN-resistant monocytic AML retains a targetable vulnerability to canonical pyroptosis. **(A)** Correlation between canonical pyroptosis signature scores and venetoclax (VEN) area under the dose-response curve (AUC) in the BeatAML cohort. Points are colored by FAB group, and Spearman correlation coefficients and *P* values are shown for all samples and the indicated FAB groups. **(B)** Correlation between canonical pyroptosis signature scores and VEN selective drug sensitivity score (sDSS) in the FIMM-AML cohort. **(C)** Kaplan-Meier overall survival curves of AML patients receiving VEN-based therapy in the BeatAML cohort, stratified into high- and low-score groups by the median canonical pyroptosis signature score. **(D-F)** Correlation between *ex vivo* responses to VEN and VBP in primary AML specimens, assessed by CCK-8 relative viability (D), relative viability by Annexin V/DAPI flow cytometry (E) and LDH-derived viability (F). **(G-I)** Correlation between *ex vivo* responses to VEN and LPS+Nig-induced canonical pyroptosis in primary AML specimens, assessed by CCK-8 relative viability (G), relative viability by Annexin V/DAPI flow cytometry (H) and LDH-derived viability (I). **(J)** Representative flow-cytometry plots of primary FAB M5 AML cells from patient AML-68 treated *ex vivo* with DMSO, VEN, VBP, VEN+VBP, or LPS+Nig. Top panels show Annexin V/DAPI staining. Bottom panels show CD45⁺CD4⁺, CD45⁺CD14⁺ and CD45⁺CD64⁺ monocytic fractions among gated viable cells defined as Annexin V⁻DAPI⁻. Numbers indicate the percentage of cells in the indicated quadrants or gates. **(K)** Quantification of the relative viability of primary AML cells following the indicated *ex vivo*treatments, corresponding to the Annexin V/DAPI flow-cytometry assay illustrated in (J). **(L)** Quantification of CD45⁺CD4⁺, CD45⁺CD14⁺ and CD45⁺CD64⁺ fractions among viable primary AML cells following the indicated *ex vivo* treatments, corresponding to the marker-gating strategy illustrated in (J). *Ex vivo* drug treatments were performed as described in Figure 3; in combination experiments, cells were exposed to VEN for 36 h, with VBP added during the final 24 h. In (A) and (B), each dot represents one AML sample; in (D-I), each dot represents one primary AML specimen. Fitted lines are shown in (A-B) and (D-I); shaded bands in (A) and (B) indicate 95% confidence intervals. Spearman correlation coefficients and *P* values are indicated in the plots. In (C), numbers at risk are shown below the plot, and the *P* value was calculated using the log-rank test. In (D-I), CCK-8 and flow-cytometry relative viability values were normalized to matched DMSO-treated controls. For LDH assays, cell death was calculated from baseline-corrected LDH release as a percentage of maximal release, and LDH-derived viability was calculated as 100% minus the LDH-derived cell death percentage. In (K) and (L), each dot represents one primary AML specimen (*n* = 20, comprising 15 M5 and 5 M4 specimens); boxes indicate the median and interquartile range, and whiskers indicate the minimum and maximum values. Values were normalized to matched DMSO controls. In (K), comparisons with DMSO were performed against the DMSO-normalized value of 100% using one-sample Wilcoxon signed-rank tests with Bonferroni correction; selected pairwise comparisons among non-DMSO groups were performed using the Friedman test followed by Dunn’s multiple-comparisons test. In (L), statistical significance was determined by repeated-measures two-way ANOVA followed by Dunnett’s multiple-comparisons test against matched DMSO controls within each marker population. ns, not significant; \**P* < .05, \*\**P* < .01, \*\*\**P* < .001, \*\*\*\**P* < .0001.

We next analyzed *ex vivo* drug-sensitivity profiles of primary AML samples. Consistent with previous studies,^15,19,32–34^ M4/M5 samples were more resistant to VEN and more sensitive to VBP than M0-M3 samples (Figure 3A). Of note, across all AML samples, regardless of FAB subtype, VEN and VBP responses correlated inversely (Figure 4D-F). LPS+Nig produced similar results (Figure 4G-I), further supporting canonical pyroptosis susceptibility in VEN-resistant AML. After FAB stratification, this reciprocal pattern remained evident in combined M4/M5 and M5-only samples (supplemental Figure 10A-B; supplemental Figure 11A-B). Similar trends in combined M0-M3 AML, M4-only AML, and even ALL samples were observed but did not reach significance (supplemental Figure 10C-E; supplemental Figure 11C-E), likely because of low response rates to VBP or LPS+Nig in non-monocytic leukemia and limited sample sizes. Consistently, VEN-resistant samples were more sensitive to both VBP and LPS+Nig than VEN-sensitive samples (supplemental Figure 12A-F).

VEN-based regimens have been reported to spare and enrich monocytic AML populations, driving refractory and relapsed disease.^15,24,33,34,37^ In primary samples containing both primitive and monocytic populations, VEN preferentially depleted non-monocytic primitive cells, increasing the proportion of surviving cells expressing monocytic markers such as CD4, CD14, and CD64 (Figure 4J, L). Notably, these VEN-resistant monocytic cells were precisely the population sensitive to VBP and LPS+Nig, and combining VBP with VEN counteracted the reciprocal population enrichment caused by either agent alone (Figure 4J, L), resulting in deeper AML-cell eradication (Figure 4K).

We further generated a VEN-resistant subline of MONO-MAC-6 cells (supplemental Figure 13A), which is monocytic but relatively less VBP-sensitive (Figure 2A, C, D), to determine whether acquired VEN resistance can sensitize cells to VBP. The VEN-resistant cells showed increased VBP sensitivity, with an expanded Annexin V⁺/DAPI⁺ pyroptotic population and increased GSDMD cleavage relative to parental cells (supplemental Figure 13B-D). RNA-seq also showed a higher monocytic differentiation score and elevated canonical pyroptosis gene expression (supplemental Figure 13E-H). Thus, beyond the selection model of monocytic versus primitive clones observed in primary AML, acquired VEN resistance in a relatively homogeneous cell line could increase monocytic differentiation and canonical pyroptosis gene expression, directly rendering cells susceptible to pyroptosis.

Collectively, canonical pyroptosis activity and VBP sensitivity are positively associated with VEN resistance in AML. Adding canonical pyroptosis inducers such as VBP to VEN-based regimens may therefore improve therapeutic outcomes.

### 5. Val-boroPro suppresses venetoclax-resistant AML progression *in vivo*

To evaluate whether VBP overcomes VEN resistance *in vivo*, we established VEN-resistant cell line-derived xenograft (CDX) and patient-derived xenograft (PDX) models in NCG mice. The CDX used a VEN-resistant subline of MV4-11, a monocytic AML line whose parental cells are relatively VEN-sensitive (supplemental Figure 13I), and acquisition of VEN resistance did not alter its VBP sensitivity (supplemental Figure 13J). The PDX used primary FAB M5 cells from patient AML-82, which were VBP-sensitive and highly VEN-resistant (supplemental Figure 14). CDX mice were randomized 1 week after transplantation, and PDX mice were randomized after confirmed engraftment, then treated with vehicle, VEN, VBP, or VEN+VBP for 4 weeks (Figure 5A).

**Figure 5.**
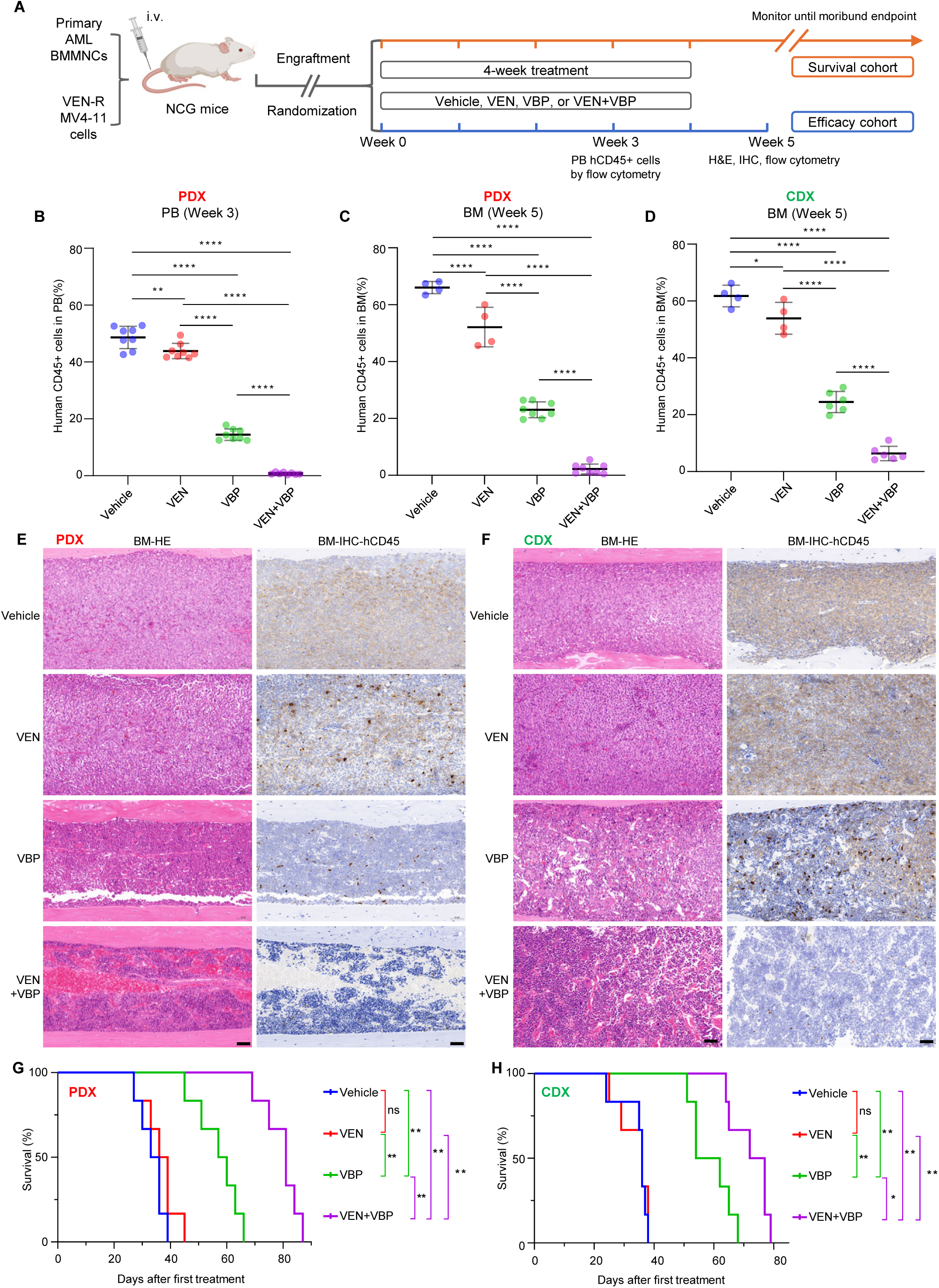
VBP suppresses VEN-resistant monocytic AML progression in CDX and PDX models. **(A)** Schematic of the *in vivo* treatment strategy for CDX and PDX models. VEN-resistant MV4-11 cells or primary AML BMMNCs from patient AML-82 (FAB M5) were transplanted intravenously into NCG mice. CDX mice were randomized one week after transplantation, whereas PDX mice were randomized after leukemic engraftment. Mice then received vehicle, VEN, VBP, or VEN+VBP for 4 weeks. Peripheral blood leukemic burden was assessed by flow-cytometric quantification of hCD45⁺ cells at treatment week 3, followed by flow-cytometric, histological, and immunohistochemical analyses at week 5 in the efficacy cohort, or survival monitoring until predefined humane endpoints in the survival cohort. **(B-D)** Flow-cytometric quantification of hCD45⁺ leukemic cells in peripheral blood from PDX mice after 3 weeks of treatment (B) and in bone marrow from PDX (C) and CDX (D) mice at week 5. **(E-F)** Representative H&E staining and hCD45 immunohistochemistry of BM sections from PDX (E) and CDX (F) mice at week 5 in the indicated treatment groups. Scale bars, 50 μm. **(G-H)** Kaplan-Meier survival curves of PDX (G) and CDX (H) mice treated with vehicle, VEN, VBP, or VEN+VBP. In (B-D), each dot represents one mouse, and data are shown as mean ± s.d. PDX groups were initially assigned *n* = 8 mice each, whereas CDX groups were initially assigned *n* = 6 mice each; the numbers analyzed at week 5 varied because mice reaching humane endpoints before the scheduled terminal collection were unavailable for terminal analyses. Statistical significance was determined by one-way ANOVA followed by Tukey’s multiple-comparisons test for indicated comparisons. Survival analyses in (G-H) included *n* = 6 mice per group, and pairwise survival comparisons were performed using the log-rank (Mantel-Cox) test with Holm-Šídák correction for multiple comparisons. ns, not significant; \**P* < .05, \*\**P* < .01, \*\*\*\**P* < .0001.

On day 21 of treatment, when the first vehicle- and VEN-treated CDX mice in the cohorts for efficacy analysis became moribund (Figure 5A), peripheral blood (PB) was collected from both models. VBP markedly reduced PB hCD45^+^ cells in both models, whereas VEN did not significantly reduce CDX leukemic burden and had limited activity in the PDX (Figure 5B; supplemental Figure 15A). VEN+VBP produced the greatest reduction, leaving PB hCD45^+^ cells nearly undetectable by flow cytometry (Figure 5B; supplemental Figure 15A). At week 5, 1 week after treatment ended, PB and BM hCD45^+^ cells were markedly lower with VBP than VEN and were further reduced by VEN+VBP in both models (Figure 5C-D; supplemental Figure 15B-C). Consistently, week 5 H&E staining and hCD45 immunohistochemistry showed that VBP attenuated the extensive leukemic infiltration in the BM, spleen, and liver, and VEN+VBP largely suppressed leukemic growth and preserved organ architecture (Figure 5E-F; supplemental Figure 15D-E). In parallel cohorts for survival analysis, VEN did not significantly prolong survival versus vehicle, whereas VBP prolonged survival versus vehicle and VEN, and VEN+VBP further extended survival beyond either monotherapy in both models (Figure 5G-H). Thus, VBP suppresses VEN-resistant AML progression *in vivo*, with additional benefit from combining it with VEN.

Collectively, our findings strongly support clinical evaluation of VBP alone or with VEN-based regimens in monocytic AML, AML with minor monocytic clones, and AML harboring mutations associated with monocytic differentiation and VEN resistance.

## Discussion

Evasion of oncogene-induced apoptosis enables malignant transformation, and apoptosis remains the principal programmed cell death engaged by radiation and chemotherapy.^44,58^ By directly countering this cancer hallmark, VEN has reshaped AML treatment, yet resistance is very common.

Phenotypic states of cancer are more plastic and harder to assess rigorously than genetic alterations,^59^ perhaps explaining why monocytic AML is absent from ELN2024 risk stratification^25^ despite predicting poor outcomes with VEN-based therapy.^33,35,36^ Given AML’s rapid course, selecting pre-existing monocytic subclones^33^ with monocytic-associated mutations^14,38–40^ is faster than acquiring *de novo BAX*^26^ or *BCL2* (frequent in VEN-treated CLL^60^) mutations to confer resistance.^14^ Indeed, *RAS*-mutant clones drive VEN-resistance and relapse with monocytic features.^24^ In brief, multiple VEN-resistance drivers exploit monocytic AML’s MCL1 rather than BCL2 dependence. Here, we show that the driver-induced monocytic state retains canonical pyroptotic capacity that can itself be targeted to overcome VEN resistance.

Recently, pyroptosis has emerged as the best-characterized cell-death program after apoptosis. It can directly kill cancer cells and trigger antitumor immunity,^44^ but cancers often silence GSDME/GSDMB-mediated pyroptosis via *GSDME* promoter methylation^44^ or *GSDMB* isoform switching.^45^ Unlike apoptosis and non-GSDMD mediated pyroptosis, canonical GSDMD pyroptosis serves antimicrobial defense rather than tumor surveillance; its machinery may therefore escape negative selection during malignant evolution.^44,46^ Crosstalk with tumor-suppressive intrinsic apoptosis further distinguishes this program: apoptosis executioner caspase-3 cleaves and activates GSDME/B^46^ but cleaves GSDMD to abolish its pore-forming capacity.^61^ Consistent with this idea, incompletely differentiated monocytic AML retained canonical pyroptosis machinery at levels comparable to normal monocytes. Because *IL1B*/*IL18*-mediated inflammation is dispensable for lytic execution, we excluded these genes from our canonical pyroptosis signature.

Accordingly, monocytic AML is the principal subtype susceptible to LPS+Nig and VBP across cell lines and primary samples. We also noticed that most previously reported VBP-sensitive AML lines were FAB M5.^48^ With VBP having entered clinical evaluation,^49^ our findings support prospective patient enrichment or stratification by monocytic differentiation.

Most solid-tumor cells are highly resistant to BH3 mimetics targeting a single anti-apoptotic BCL2 protein because efficient apoptosis requires combined BCL-XL and MCL1 inhibition.^44^ However, on-target toxicities constrain MCL1 or BCL-XL inhibition,^14,46^ leaving selective BCL2 inhibitors as the only approved anticancer agents directly harnessing apoptosis, with indications confined to hematopoietic malignancies. This constraint also limits options after VEN failure.^14^ Our data offer an alternative: canonical pyroptosis gene expression and VBP sensitivity correlated positively with VEN resistance across FAB M0-M5, encompassing most AML cases; VBP and LPS+Nig eliminated VEN-persistent monocytic cells; VBP prolonged survival in VEN-resistant xenografts. Adding VBP to VEN-based regimens could therefore target pre-existing and treatment-enriched monocytic populations. If approved, VBP would establish pyroptosis as the second cell-death program directly harnessed by an approved cancer therapy after apoptosis, again in hematopoietic malignancies.

VBP activates human NLRP1 by binding DPP9, disrupting DPP9-mediated sequestration of NLRP1-CT, and promoting proteasomal degradation of repressive NLRP1-NT; freed NLRP1-CT then activates caspase-1 in an ASC-dependent manner.^53,62^ VBP-induced CARD8 inflammasome activation probably follows a similar pattern, except that CARD8 itself activates caspase-1 independent of ASC.^48,53^ Because *NLRP1*, *ASC* and *CARD8* are all expressed in monocytic AML, whether both inflammasomes contribute to VBP-induced pyroptosis remains unresolved. Pyroptosis releases DAMPs and tumor antigens. Whether these DAMPs engage the DAMP-sensing PRRs expressed in monocytic AML to amplify the initial VBP-induced pyroptosis, and whether released tumor antigens promote DC presentation and cross-prime CD8^+^ T cells,^44^ to mediate secondary AML killing, warrant further investigation.

This work reveals that VEN-resistant and monocytic AML are highly primed for canonical pyroptosis. Once the trial establishes VBP safety, the VEN+VBP combination merits clinical evaluation.

## Acknowledgments

This study was supported by the Provincial Natural Science Foundation of Hunan (2023JJ20023 and 2025JJ81126), the National Natural Science Foundation of China (82200198 and 82370177), and the Municipal Science and Technology Innovation Project of Xiangtan (SFYB20240011).

## Authorship Contributions

Conceptualization: G. Zhu. Original draft preparation: G. Zhu, Q. Wang, S. Xu. Review and editing: G. Zhu, X. Yang, H. Yang, Q. Wang, S. Xu. Experiments, data collection and analysis: Q. Wang, S. Xu, J. Cai, Q. Peng, S. Cao, Y. Zhang, P. Lu, X. Guo, L. Liu, W. Fu, S. Shao, J. Zhu, B. Qing, H. Li, K. Liu. Supervision: G. Zhu, X. Yang, H. Yang. Funding acquisition: G. Zhu, X. Yang, K. Zhu. All authors have read and agreed to the final version of the manuscript.

## Data Sharing Statement

Publicly available microarray, bulk RNA-seq, scRNA-seq, clinical annotation, and drug-response datasets analyzed in this study were obtained from TCGA, GEO, CCLE, ArrayExpress, and published studies; accession numbers, data types, and source references are provided in supplemental Table 1. Raw cell-line RNA-sequencing reads generated in this study are publicly available in the NCBI Sequence Read Archive (SRA) under BioProject accession PRJNA1516960. Processed RNA-sequencing data have been submitted to GEO, and the GEO accession number will be provided once assigned. Curated pyroptosis-related gene signatures are provided in supplemental Table 2. Additional original data are available from the corresponding author upon reasonable request.

