## Supplemental Material for "Venetoclax Resistance Creates Canonical Pyroptosis Vulnerability in AML"

#### **Supplemental Methods**

##### **Reagents and antibodies**

Reagents used in this study included Val-boroPro mesylate (VBP; AmBeed, Cat#A264157), venetoclax (VEN; Selleck, Cat#S8048), lipopolysaccharide (LPS; MCE, Cat#HY-D1056), nigericin (MCE, Cat#HY-127019), CCK-8 reagent (APExBIO Technology, Cat#K1018), and LDH Cytotoxicity Assay Kit (Beyotime Biotechnology, Cat#C0017).

Flow-cytometric reagents and antibodies used for primary leukemia specimens included Annexin V-FITC (Elabscience, Cat#E-CK-A111), DAPI (Elabscience, Cat#E-CK-A163), anti-human CD4 (BioLegend, Cat#300552, APC, clone RPA-T4), anti-human CD14 (BioLegend, Cat#301824, PerCP/Cyanine5.5, clone M5E2), anti-human CD45 (BioLegend, Cat#304058, PE, clone HI30), and anti-human CD64 (BioLegend, Cat#305022, PE/Cyanine7, clone 10.1). For mouse xenograft analyses, the anti-human CD45 antibody described above was used to identify hCD45<sup>+</sup> leukemic cells, and anti-mouse CD45.2 (BioLegend, Cat#109827, PerCP/Cyanine5.5, clone 104) was additionally used when required to distinguish murine hematopoietic cells from human leukemic cells.

##### **Cell culture**

Human leukemia cell lines MOLM-16, K-562, KG-1, KG-1a, HL-60, Kasumi-1, NB4, OCI-AML2, OCI-AML3, EOL-1, MONO-MAC-6, MOLM-13, MV4-11, NOMO-1, THP-1, U-937, HEL, TF-1, CMK, MEG-01, BALL-1, NALM-6, REH, and MOLT-4 were obtained from Procell and authenticated by short tandem repeat (STR) genotyping. Cells were cultured in RPMI 1640 medium (Procell, Cat#PM150110) supplemented with 10% fetal bovine serum (FBS; Yeasen, Cat#40132ES) and 1× penicillin/streptomycin (Biosharp, Cat#BL505A). TF-1 cells were additionally supplemented with 2 ng/mL granulocyte-macrophage colony-stimulating factor (GM-CSF). All cells were maintained at 37 °C in a humidified incubator with 5% CO<sub>2</sub>.

#### **Generation of VEN-resistant cell lines**

VEN-resistant MONO-MAC-6 and MV4-11 cells were generated by long-term culture with stepwise increases in VEN concentration. Resistant cells were maintained in medium containing 10 μM VEN. Before experiments, VEN-resistant cells were cultured in VEN-free medium for at least 2 weeks. Parental cells were maintained in parallel under standard culture conditions.

#### **Drug treatments**

VBP and VEN were dissolved in DMSO, LPS was dissolved in sterile water, and nigericin was dissolved in absolute ethanol. Stock solutions were aliquoted and stored according to the manufacturers' instructions. DMSO-treated cells were used as the reference control for normalized analyses, including experiments involving LPS and Nig.

For dose-response assays in leukemia cell lines, cells were treated with increasing concentrations of VBP or VEN for 48 h. For representative flow-cytometric analyses and LDH release assays in leukemia cell lines, cells were treated with VBP at 0.5, 5 and 50 μM for 48 h. For primary leukemia *ex vivo* studies, VEN was used at 0.5 μM for 36 h and VBP at 5 μM for 24 h unless otherwise indicated. In combination-treatment experiments, cells were exposed to VEN for 36 h, with VBP added during the final 24 h of treatment. For LPS plus nigericin (LPS+Nig) treatment, leukemia cell lines and primary leukemia cells were primed with LPS at 10 μg/mL for 24 h, followed by nigericin treatment at 10 μM for 1 h.

#### **CCK-8 cell viability assays**

Cell viability was measured using the CCK-8 assay. All leukemia cell lines used in these assays were suspension cells. For cell-line assays, cells were seeded at $1 \times 10^4$ cells per well in 96-well plates in 100 μL complete medium and equilibrated overnight before drug treatment. For primary leukemia specimens, thawed cells were seeded at $5 \times 10^4$ cells per well in 96-well plates in 100 μL medium and treated as indicated, with each condition performed in triplicate when sufficient cells were available. After treatment, 10 μL CCK-8 reagent (APExBIO Technology, Cat#K1018) was added to each well, and plates were incubated for 4 h at 37 °C. Absorbance was measured at 450 nm using a microplate reader. Relative viability was normalized to the corresponding DMSO-treated controls, and CCK-8-derived cell death was calculated as 100% minus relative viability. Dose-response curves were fitted using a three-parameter logistic model in GraphPad Prism 10.

##### **Flow cytometry**

For flow-cytometric analyses, cell viability and cell death were assessed by Annexin V/DAPI staining. After drug treatment, cells were washed with 1× rinse buffer and stained with Annexin V-FITC and DAPI according to the manufacturer's instructions. Viable cells were defined as Annexin V<sup>-</sup>DAPI<sup>-</sup>. Relative viability was normalized to the corresponding DMSO-treated controls, and cell death was calculated as 100% minus relative viability or as the percentage of Annexin V<sup>+</sup> and/or DAPI<sup>+</sup> cells, as indicated.

For primary leukemia immunophenotyping, thawed primary leukemia BMMNCs were seeded at $5 \times 10^5$ cells per well in 48-well plates in 500 μL medium. After treatment, cells were stained with anti-human CD4 (BioLegend, Cat#300552, APC, clone RPA-T4), anti-human CD14 (BioLegend, Cat#301824, PerCP/Cyanine5.5, clone M5E2), anti-human CD45 (BioLegend, Cat#304058, PE, clone HI30) and anti-human CD64 (BioLegend, Cat#305022, PE/Cyanine7, clone 10.1), together with Annexin V-FITC (Elabscience, Cat#E-CK-A111) and DAPI (Elabscience, Cat#E-CK-A163). Samples were analyzed on a Beckman Coulter CytoFLEX flow cytometer, and data were processed using FlowJo v10.9.0.

##### **LDH release assay**

LDH release was measured using the LDH Cytotoxicity Assay Kit (Beyotime Biotechnology, Cat#C0017) according to the manufacturer's instructions. Cells were seeded at $5 \times 10^4$ cells per well in 48-well plates for cell-line assays, and supernatants from primary leukemia cell cultures were collected from the corresponding 48-well *ex vivo* cytotoxicity assays. Maximal LDH release was induced using the kit-provided LDH release reagent at 10% of the culture volume. Absorbance of the clarified supernatants was measured at 490 nm using a microplate reader. LDH release was calculated as the percentage of maximal release after subtraction of the corresponding DMSO-treated control baseline. Where indicated, LDH-derived viability was calculated as 100% minus the LDH-derived cell death percentage.

##### **Immunoblotting**

Cells were seeded at $1 \times 10^6$ cells per well in 6-well plates and treated with DMSO or the indicated compounds. OCI-AML2, OCI-AML3, EOL-1, MOLM-13, NOMO-1, and THP-1 were treated with VBP at 10 μM for 24 h. MV4-11, MONO-MAC-6, MOLM-16, K-562, KG-1, KG-1a, HL-60, Kasumi-1, NB4, U-937, HEL, TF-1, CMK, MEG-01, BALL-1, NALM-6, REH and MOLT-4 were treated with VBP at 10 μM for 48 h. For LPS plus nigericin treatment, cells were primed with LPS at 10 μg/mL for 24 h, followed by nigericin treatment at 10 μM for 1 h.

After treatment, cells were collected and washed with cold PBS. Cell pellets were lysed in SDS lysis buffer (Beyotime, Cat#P0013G). Protein concentrations were determined using the BCA assay (Thermo Fisher Scientific, Cat#23225). Proteins were separated on 12% SDS-PAGE gels and transferred onto Immobilon PVDF membranes (Sigma-Aldrich, Cat#IPVH00010). Membranes were blocked with 5% non-fat milk in TBST for 1 h and incubated with primary antibodies overnight at 4 °C. Membranes were then incubated with HRP-conjugated secondary antibodies corresponding to the host species of the primary antibodies. HRP signals were detected using a Bio-Rad ChemiDoc imaging system.

Antibodies for immunoblotting were as follows: CARD8 (Santa Cruz Biotechnology, Cat#sc-81213, 1:1,000), caspase-1 (Abcam, Cat#ab179515, 1:1,000), GSDMD (Abcam, Cat#ab210070, 1:1,000), cleaved N-terminal GSDMD (GSDMD-N; Abcam, Cat#ab215203, 1:1,000), β-actin (CST, Cat#3700, 1:5,000), anti-rabbit IgG H&L (HRP) (Abcam, Cat#ab6721, 1:5,000), and anti-mouse IgG H&L (HRP) (Abcam, Cat#ab6728, 1:5,000).

##### **Primary leukemia specimens and *ex vivo* culture**

Primary bone marrow specimens from 106 patients with acute myeloid leukemia

(AML) and 36 patients with acute lymphoblastic leukemia (ALL) were obtained at Xiangya Hospital, Central South University. Written informed consent was obtained from all participants before sample collection. Clinical and molecular characteristics of the primary leukemia specimens are summarized in supplemental Table 3.

Bone marrow aspirates were collected from enrolled patients, and bone marrow mononuclear cells (BMMNCs) were isolated by Ficoll-Paque density-gradient centrifugation (TBD Science, Cat#LTS1077) according to the manufacturer's instructions. Cells were washed with 1× rinse buffer consisting of PBS supplemented with BSA and EDTA. Residual erythrocytes were lysed with ACK lysis buffer, and cells were counted and cryopreserved until use. Upon thawing, primary leukemia BMMNCs were cultured in RPMI 1640 medium supplemented with 10% FBS, 1× penicillin/streptomycin and 0.1 mM 2-mercaptoethanol.

##### **Xenograft studies**

All animal experiments were performed in the animal facility of Hunan University and approved by the Animal Care and Welfare Committee of Hunan University. Six-week-old female NCG mice (NOD/ShiLtJGpt-*Prkdc*<sup>em26Cd52</sup>*Il2rg*<sup>em26Cd22</sup>/Gpt; GemPharmatech) were used for xenograft studies and maintained under specific pathogen-free conditions with ad libitum access to food and water. All mice received 100 cGy total-body irradiation prior to transplantation.

For the cell-line-derived xenograft (CDX) model, $1 \times 10^6$ viable VEN-resistant MV4-11 cells suspended in 200 μL PBS were injected intravenously via the tail vein. One week after transplantation, mice were randomly allocated to efficacy and survival cohorts and subsequently to treatment groups using a computer-generated random-number sequence.

For the patient-derived xenograft (PDX) model, $3 \times 10^6$ viable BMMNCs obtained from patient AML-82 (FAB M5) and suspended in 200 μL PBS were injected intravenously via the tail vein. Leukemic engraftment was monitored weekly by flow-cytometric quantification of hCD45<sup>+</sup> cells in peripheral blood. Eligibility for randomization was predefined as hCD45<sup>+</sup> cells exceeding 1% of viable (DAPI<sup>-</sup>) singlet peripheral blood cells after red blood cell lysis. At approximately 3 weeks after transplantation, eligible mice were randomly allocated to efficacy and survival cohorts. Within each cohort, mice were ranked according to baseline peripheral blood hCD45<sup>+</sup> leukemic burden and randomized in blocks of four using a computer-generated random-number sequence, with one mouse from each block assigned to each treatment group. Treatment commenced on the day of randomization.

Within each cohort, mice were assigned to one of four treatment groups: vehicle, VEN, VBP, or VEN+VBP. VEN was freshly prepared in 10% DMSO, 40% PEG300, 5% Tween-80, and 45% saline and administered by oral gavage at 100 mg/kg. VBP was dissolved in PBS and administered intraperitoneally at 2 mg/kg. Vehicle-treated mice received the VEN vehicle by oral gavage and 100 μL PBS intraperitoneally. Each monotherapy group received the corresponding drug and the matched vehicle for the other agent, whereas the combination group received both drugs. Treatments and corresponding vehicles were administered once daily, 5 days per week, for 4 weeks.

Leukemic burden in peripheral blood and bone marrow was quantified by flow cytometry as the percentage of hCD45<sup>+</sup> cells among viable (DAPI<sup>-</sup>) singlet cells in the corresponding compartment following red blood cell lysis. In the efficacy cohorts, peripheral blood was collected at treatment week 3. At the terminal time point, one week after completion of the 4-week treatment period, peripheral blood was collected from the lateral tail vein before mice were euthanized by CO<sub>2</sub> inhalation. Bone marrow, spleen, and liver were subsequently harvested for downstream analyses.

All mice were monitored daily and immediately euthanized by CO<sub>2</sub> inhalation upon reaching any of the following predefined humane endpoints: a loss of >20% of body weight relative to that at treatment initiation, labored breathing, inability to access food or water, severe hindlimb weakness or paralysis impairing ambulation, minimal responsiveness to external stimuli, or a moribund state. In the efficacy cohorts, mice reaching a humane endpoint before the scheduled terminal collection were euthanized and were therefore unavailable for terminal tissue analyses; tissues were collected from all mice remaining at the scheduled endpoint. In the survival cohorts, survival was defined as the interval from treatment initiation to the humane endpoint, with euthanasia at the endpoint recorded as an event. No animals or data points were otherwise excluded after randomization.

##### **Histology and immunohistochemistry**

Liver, spleen and bone marrow tissues were collected from mice remaining in the efficacy cohorts at the scheduled terminal endpoint, fixed in 4% paraformaldehyde, embedded in paraffin and sectioned. Bone marrow samples were decalcified in EDTA before embedding. Bone marrow sections were cut at 3 μm, and spleen and liver sections at 2 μm. Hematoxylin and eosin (H&E) staining was performed to assess tissue architecture and leukemic infiltration.

Immunohistochemistry for hCD45 was performed on bone marrow sections after antigen retrieval in citrate buffer (pH 6.0), using anti-human CD45 antibody (Abcam, Cat#ab208022, 1:400), HRP-conjugated goat anti-rabbit secondary antibody (Abcam, Cat#ab205718, 1:2,000) and DAB visualization. Images were acquired using Panoramic MIDI or Panoramic 250 FLASH digital slide scanners (3DHISTECH). Scale bars represent 50 μm.

##### **Public datasets and clinical cohorts**

Pan-cancer analyses of primary tumors and cancer cell lines were performed using The Cancer Genome Atlas (TCGA) and the Cancer Cell Line Encyclopedia (CCLE), respectively. Publicly available clinical annotations, transcriptomic profiles and drug-response data were collected from TCGA, the Gene Expression Omnibus (GEO), ArrayExpress and published studies. AML-specific datasets included BeatAML, TARGET-AML, TCGA-LAML, FIMM-AML, Leucegene (GSE232129), GSE146173, GSE165656, GSE12417, GSE37642, GSE10358, GSE14468, HOVON (E-MTAB-3444), GSE132511, GSE232559, GSE185381, GSE235063 and GSE239721. Accession numbers, data types and source references are summarized in supplemental Table 1.

##### **Bulk transcriptomic analysis and batch correction**

RNA-seq expression matrices were converted to fragments per kilobase of transcript per million mapped fragments (FPKM) and log<sub>2</sub>-transformed as log<sub>2</sub>(FPKM

+ 1). For meta-analysis of RNA-seq datasets, TCGA-LAML, FIMM-AML, Leucegene (GSE232129), GSE146173 and GSE165656 were integrated and batch-corrected using the ComBat function in the sva package v3.50.0, resulting in a bulk RNA-seq meta-cohort of 794 samples.

For probe-based microarray datasets, probes were mapped to gene symbols according to the corresponding platform annotations; when multiple probes mapped to the same gene, their expression values were averaged for downstream analyses. Microarray expression matrices were normalized using the normalizeBetweenArrays function in limma v3.58.1. Datasets generated on the GPL96 platform, including GSE12417 and GSE37642, and datasets generated on the GPL570 platform, including GSE10358, GSE12417, GSE14468, GSE37642 and HOVON/E-MTAB-3444, were processed separately and batch-corrected using the removeBatchEffect function in limma v3.58.1, resulting in two platform-specific microarray meta-cohorts of 564 and 1,533 samples, respectively. Principal-component analysis (PCA) was performed before and after batch correction to assess residual batch effects.

##### **Single-cell RNA-seq analysis**

Single-cell RNA-sequencing (scRNA-seq) data from GSE232559 were used as provided by the original study, which included prior quality control and normalization. The dataset included 27 samples from 25 AML patients and 32,319 cells retained for downstream analysis. Using Seurat v4.4.0, the top 3,000 highly variable genes were selected for PCA, and the first 30 PCs were used for clustering and UMAP visualization. Sample-level batch effects were corrected using Harmony v1.2.0, and clusters were identified with FindNeighbors and FindClusters at a resolution of 5. Major cell types were annotated using clustifyr v1.21.0 with a published normal/leukemic bone marrow single-cell reference atlas.<sup>1</sup> AML cells were identified within myeloid compartments using a published Random Forest classifier (randomForest v4.7.1.1) based on an AML hierarchy framework.<sup>2</sup> AML differentiation-state groups, including Prim, Mono, Uni-MMP and Multi-MMP, were defined as described in the original study.

We additionally integrated three scRNA-seq datasets from GEO (GSE185381,

GSE235063, and GSE239721). Doublets were predicted and removed using Scrublet (v0.2.3) in a Python 3.9 environment, and cells were retained if they had >200 detected genes, 500-80,000 UMI counts, <10% mitochondrial UMIs and <5% *HBB*-derived UMIs. The final meta-cohort included 567,474 cells from 10 healthy samples and 137 AML samples retained after quality control. After normalization with Seurat, the top 3,000 highly variable genes and the first 30 PCs were used for Harmony-based integration and UMAP visualization. Cell types were annotated using the Random Forest classifier.

##### **Gene signatures and signature scoring**

Pyroptosis-related gene signatures were curated based on published reviews of pyroptosis,<sup>3</sup> as well as the Molecular Signatures Database (MSigDB, v2024.1.Hs). The curated pyroptosis-related signatures included canonical pyroptosis, non-canonical pyroptosis and other-pyroptosis signatures, together with the GOBP\_PYROPTOSIS gene set, as appropriate for each analysis. The full list of curated pyroptosis-related genes is provided in supplemental Table 2. Additional lineage and differentiation gene signatures were obtained from the LM22 immune signature matrix (CIBERSORT\_Monocytes) and the MSigDB database (BIOCARTA\_GRANULOCYTES\_PATHWAY, GAL\_LEUKEMIC\_STEM\_CELL\_UP, and GOBP\_MONONUCLEAR\_CELL\_DIFFERENTIATION), and were used as indicated.

For bulk transcriptomic profiles, signature scores were quantified using gene set variation analysis (GSVA, v1.50.5) from log-transformed normalized expression matrices unless otherwise stated. For scRNA-seq data, canonical pyroptosis signature scores in individual cells were calculated using the AddModuleScore function in Seurat. Heat maps of gene expression, signature scores or gene-set scores were displayed as row-wise Z-score-normalized values.

##### **Cell-line RNA-seq analysis**

For cell-line RNA-seq, $2 \times 10^6$ leukemia cells were collected for total RNA extraction using TRIzol reagent. RNA quality was assessed using a Qsep-400 system, and qualified RNA samples were subjected to mRNA-enriched, strand-specific RNA-seq library preparation using the VAHTS Universal V6 RNA-seq Library Prep Kit for MGI (NRM604-01) and sequencing on a BGI DNBSEQ-T7 platform. Briefly, mRNA was captured using mRNA capture beads, fragmented and reverse-transcribed into cDNA, followed by second-strand synthesis, end repair, 3' A-tailing, adaptor ligation, size selection, PCR amplification and library quality control. Raw sequencing data in FASTQ format were processed using fastp v0.20.1 to remove adaptor-contaminated reads and low-quality reads, with the parameters `-q 20 -l 100 -n 10`, generating clean reads for downstream analysis. Cleaned reads were aligned to the GRCh38 human reference genome using STAR v2.7.10b. The resulting BAM files were then used to generate read counts using featureCounts from Subread v2.0.3 with the gene annotation file (GENCODE v36). Differential expression analysis was performed using DESeq2 v1.42.1. Read count matrices were normalized using the median of ratios method from DESeq2, and *P* values were adjusted for multiple testing using the Benjamini-Hochberg false-discovery rate (FDR) procedure. Raw RNA-sequencing reads generated in this study are publicly available in the NCBI Sequence Read Archive (SRA) under BioProject accession PRJNA1516960. Processed read counts and associated sample metadata have been submitted to GEO; the GEO accession number will be provided once assigned.

##### **Clinical correlation and survival analyses**

Clinical correlation analyses were performed using Spearman correlation. For patient survival analyses, patients were stratified by either the median or quartiles of the canonical pyroptosis signature scores. In median-based analyses, patients were divided into high- and low-score groups using the median value. In quartile-based analyses, the top 25% and bottom 25% of patients were compared, with the middle 50% excluded. Kaplan-Meier curves were generated using survminer v0.4.9, and group differences were assessed by log-rank test.

##### **Ethical reporting**

The collection and use of primary leukemia specimens in this study were conducted in accordance with the Declaration of Helsinki and approved by the Ethics Committee of Xiangya Hospital, Central South University. Written informed consent was obtained from all participants who provided specimens. Animal experiments were reviewed and approved by the Animal Care and Welfare Committee of Hunan University and conducted in accordance with institutional and national guidelines.

##### **Statistical analysis**

Statistical analyses were performed using R v4.3.3 or GraphPad Prism 10 unless otherwise stated. Data are presented as mean ± s.d., mean ± s.e.m., or median with interquartile range, as indicated in the figure legends. For bioinformatic analyses, signature scores were compared using the Wilcoxon rank-sum test for two-group comparisons. For other two-group comparisons, statistical significance was determined using a two-sided Wilcoxon rank-sum test or a two-sided Student's *t*-test, as appropriate. For comparisons among multiple groups, the Kruskal-Wallis test, one-way ANOVA followed by Tukey's multiple-comparisons test, or repeated-measures two-way ANOVA followed by Dunnett's multiple-comparisons test was used as indicated. Clinical correlation analyses and patient survival analyses are described above. Mouse survival was analyzed using the log-rank (Mantel-Cox) test, with Holm-Šidák correction for pairwise comparisons where indicated. Heat maps were generated using ComplexHeatmap. For Figure 4K, comparisons with DMSO were performed using one-sample Wilcoxon signed-rank tests with Bonferroni correction, and selected pairwise comparisons among non-DMSO treatment groups were performed using the Friedman test followed by Dunn's multiple-comparisons test. Unless otherwise indicated, $P < .05$ was considered statistically significant.

### Supplemental Figure 1

A

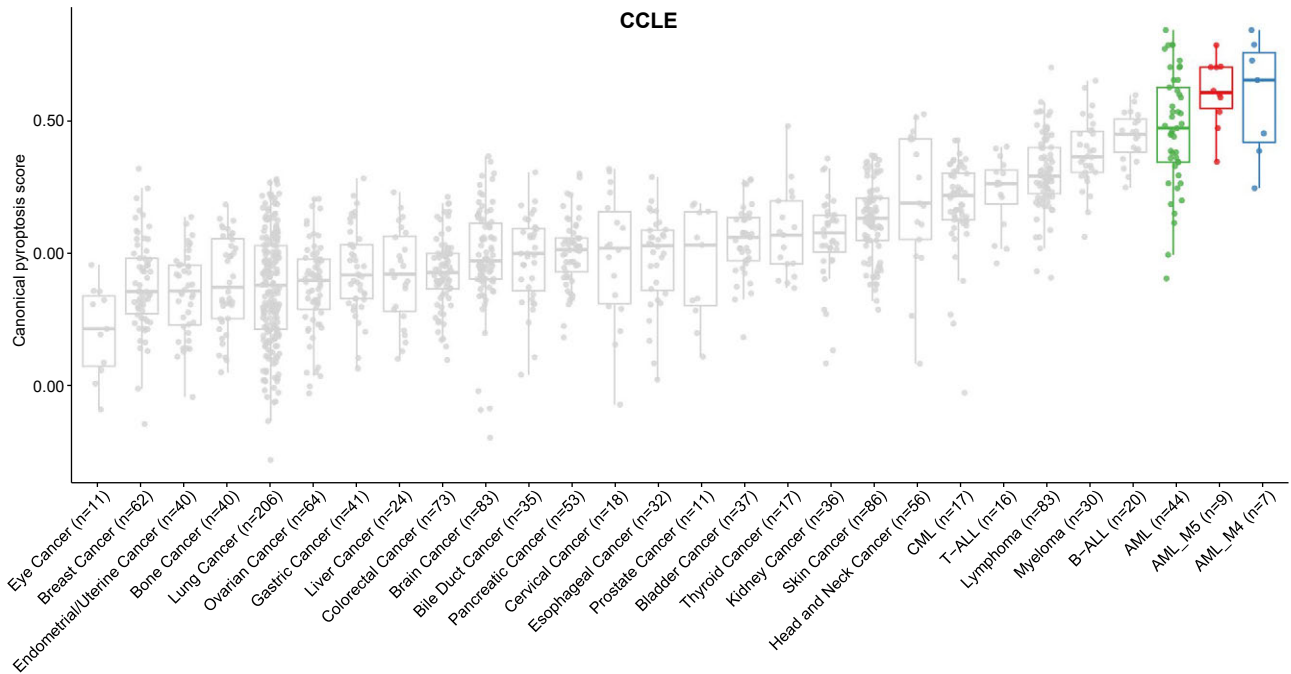

B

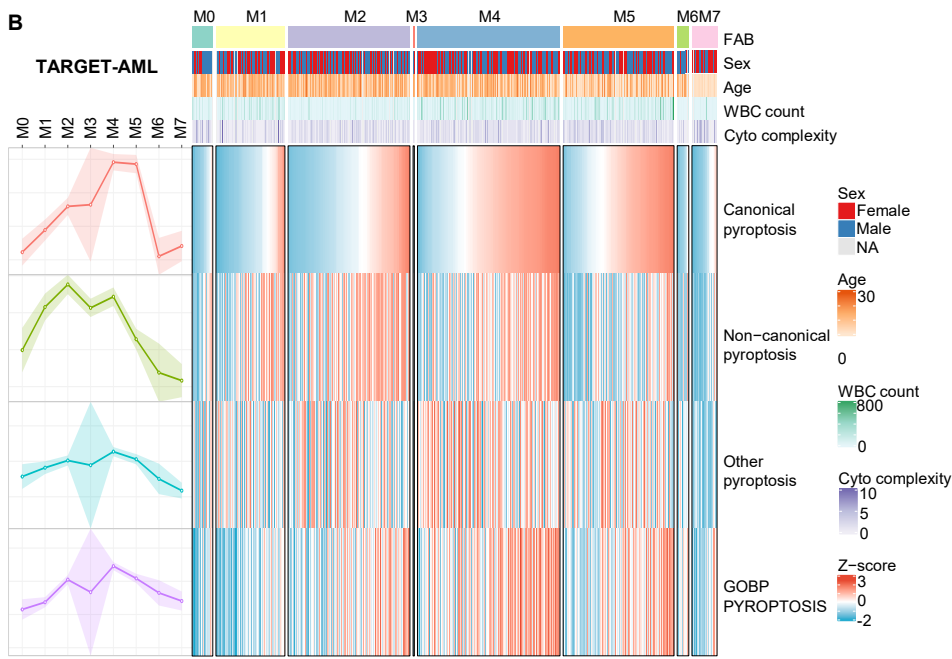

C

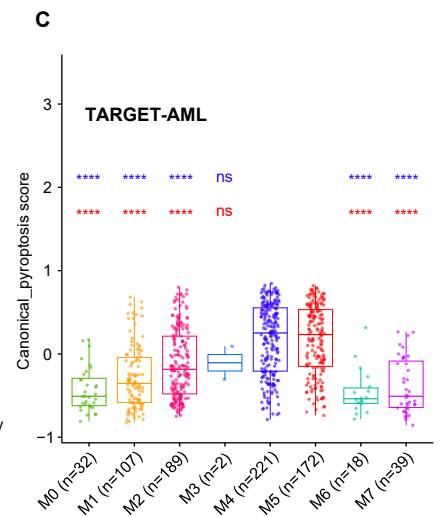

#### Supplemental Figures

##### **Supplemental Figure 1. Pan-cancer cell line and TARGET-AML analyses support enrichment of the canonical pyroptosis signature in monocytic AML.**

**(A)** Box plots showing canonical pyroptosis signature scores across cancer lineages in CCLE cell lines, with AML cell lines and their FAB M4 and M5 subsets highlighted.

**(B)** Heat map depicting the pyroptosis-related signature scores across FAB subtypes in the TARGET-AML cohort. Pyroptosis-associated genes were grouped into canonical pyroptosis, non-canonical pyroptosis, other-pyroptosis signatures, together with the GOBP\_PYROPTOSIS gene set. Samples are ordered according to FAB classification, with annotations for sex, age, white blood cell (WBC) count and cytogenetic complexity. Marginal line plots on the left summarize trends in the mean score of each pyroptosis signature across FAB subtypes. Colors represent row-wise Z-score-normalized expression values.

**(C)** Canonical pyroptosis signature scores across FAB subtypes in the TARGET-AML cohort. Violet-blue and red asterisks indicate comparisons with M4 and M5, respectively.

In (A), each dot represents one cell line. In (C), each dot represents one patient sample. Statistical significance in (C) was determined by two-sided Wilcoxon rank-sum tests for the indicated comparisons. *P* values are indicated in the plots. ns, not significant; \*\*\*\**P* < .0001.

Supplemental Figure 2

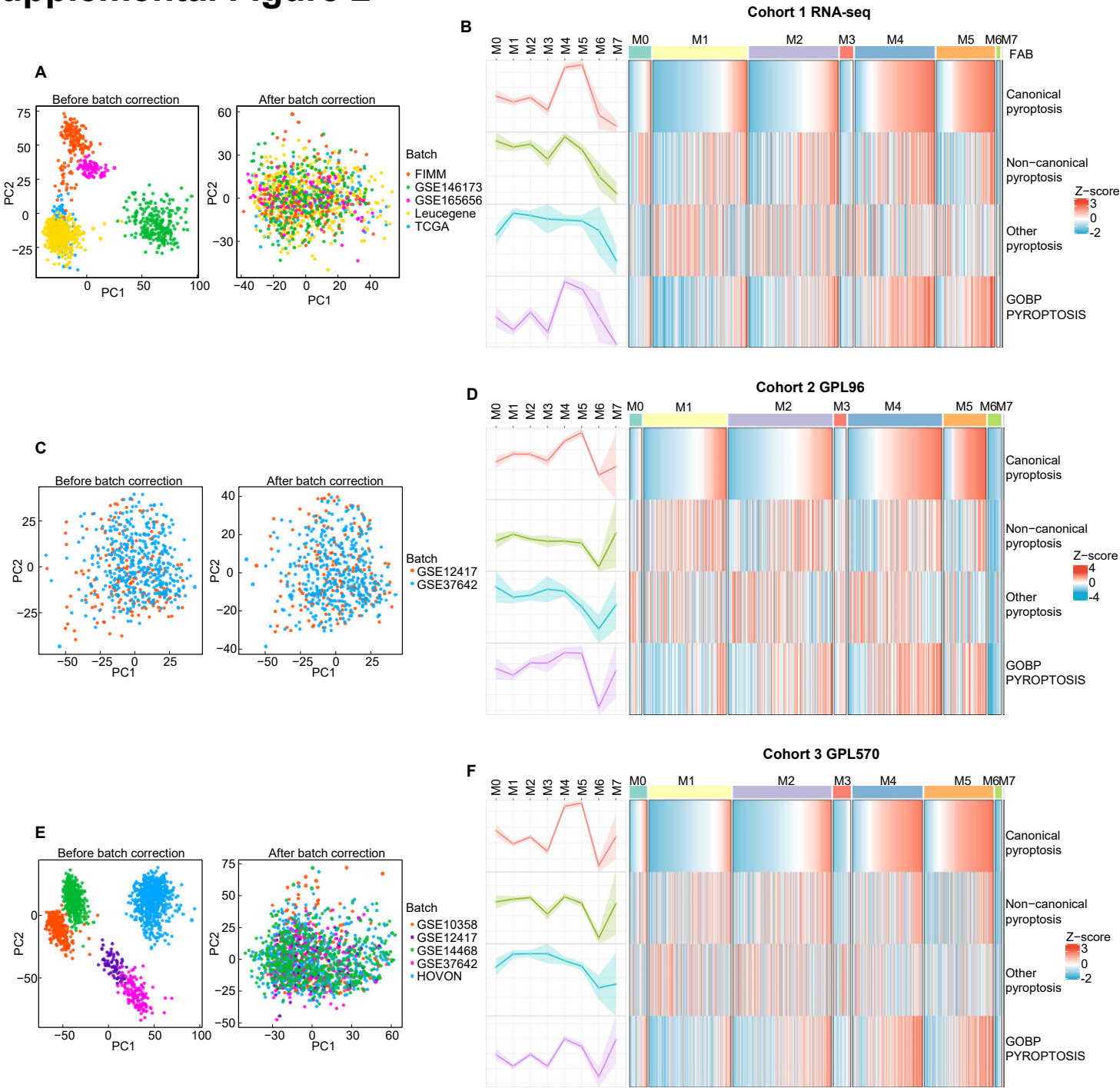

**Supplemental Figure 2. Batch-corrected AML transcriptomic cohorts validate enrichment of the canonical pyroptosis signature in monocytic AML.**

**(A,C,E)** Principal component analyses of three integrated AML transcriptomic cohorts before and after batch-effect correction. Cohort 1 comprises RNA-seq datasets from TCGA-LAML, FIMM-AML, GSE146173, GSE165656 and Leucegene **(A)**; cohort 2 comprises GPL96 microarray datasets GSE12417 and GSE37642 **(C)**; and cohort 3 comprises GPL570 microarray datasets GSE10358, GSE12417, GSE14468, GSE37642 and HOVON **(E)**. Colors denote dataset origin.

**(B,D,F)** Heat maps showing pyroptosis-related signature scores across FAB subtypes in the corresponding batch-corrected cohorts. Pyroptosis-related genes were grouped into canonical pyroptosis, non-canonical pyroptosis, other-pyroptosis signatures, together with the GOBP\_PYROPTOSIS gene set. Samples are ordered by FAB classification, and marginal line plots summarize trends in the mean score across FAB subtypes. Colors represent row-wise Z-score-normalized signature scores.

### Supplemental Figure 3

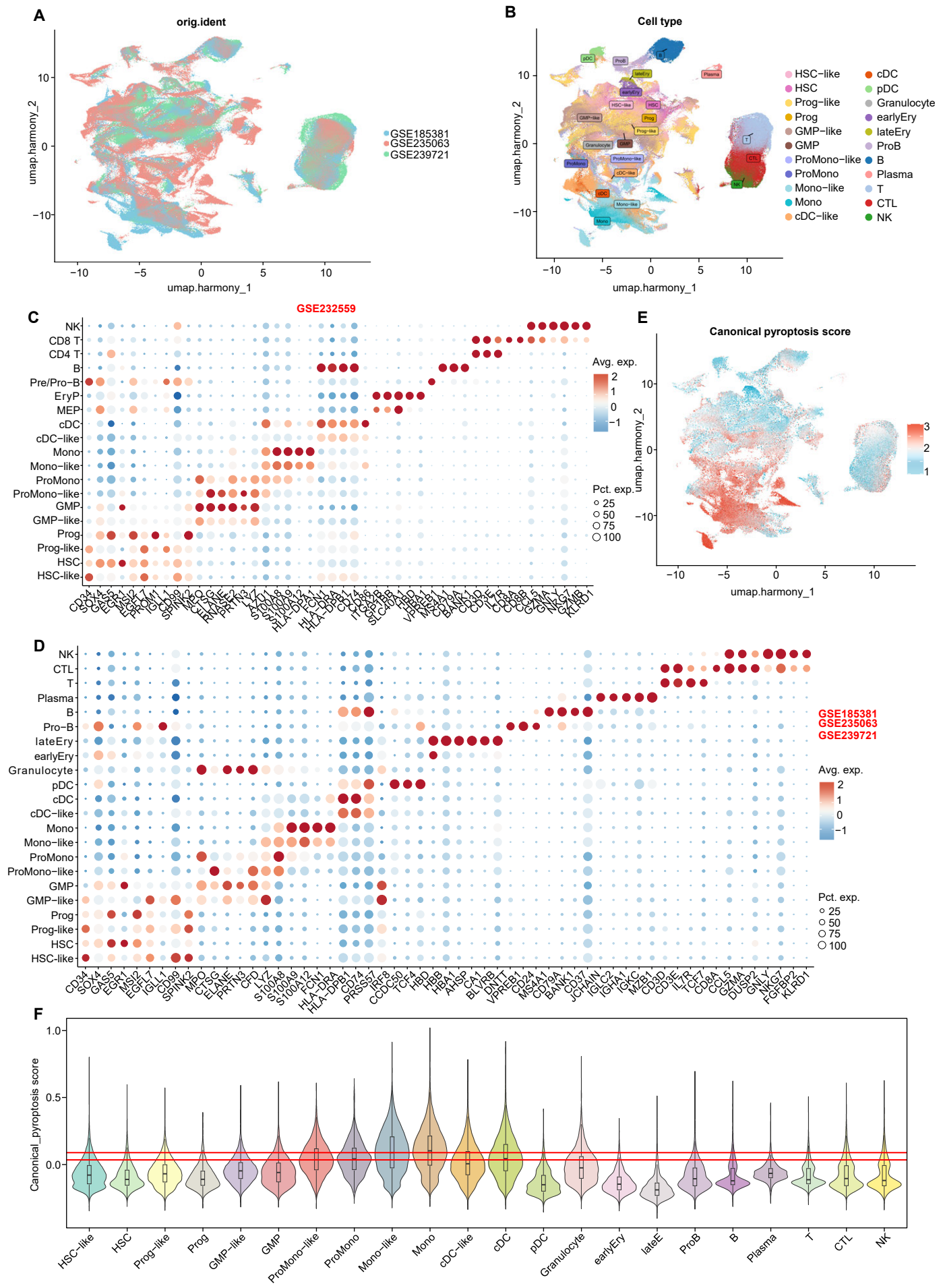

**Supplemental Figure 3. Single-cell analyses validate enrichment of the canonical pyroptosis signature in monocytic AML populations.**

**(A)** UMAP visualization of integrated single-cell transcriptomes from GSE185381, GSE235063 and GSE239721 after Harmony correction, comprising 567,474 cells from 10 healthy samples and 137 AML samples, colored by dataset of origin.

**(B)** UMAP visualization of cells in the integrated single-cell RNA-seq dataset, colored by annotated cell types identified based on canonical marker gene expression.

**(C)** Bubble plot showing representative marker-gene expression across annotated cell types in the GSE232559 dataset. Dot color indicates average expression, and dot size indicates the percentage of expressing cells.

**(D)** Bubble plot showing representative marker-gene expression across annotated cell types in the integrated GSE185381, GSE235063 and GSE239721 datasets. Dot color indicates average expression, and dot size indicates the percentage of expressing cells.

**(E)** UMAP visualization of canonical pyroptosis signature scores across all profiled cells in the integrated GSE185381, GSE235063, and GSE239721 single-cell RNA-seq dataset.

**(F)** Violin plot comparing canonical pyroptosis signature scores across annotated AML and normal hematopoietic cell populations in the integrated GSE185381, GSE235063, and GSE239721 single-cell RNA-seq dataset. The lower and upper red horizontal lines indicate the median canonical pyroptosis signature scores of the ProMono-like and Mono-like populations, respectively.

### Supplemental Figure 4

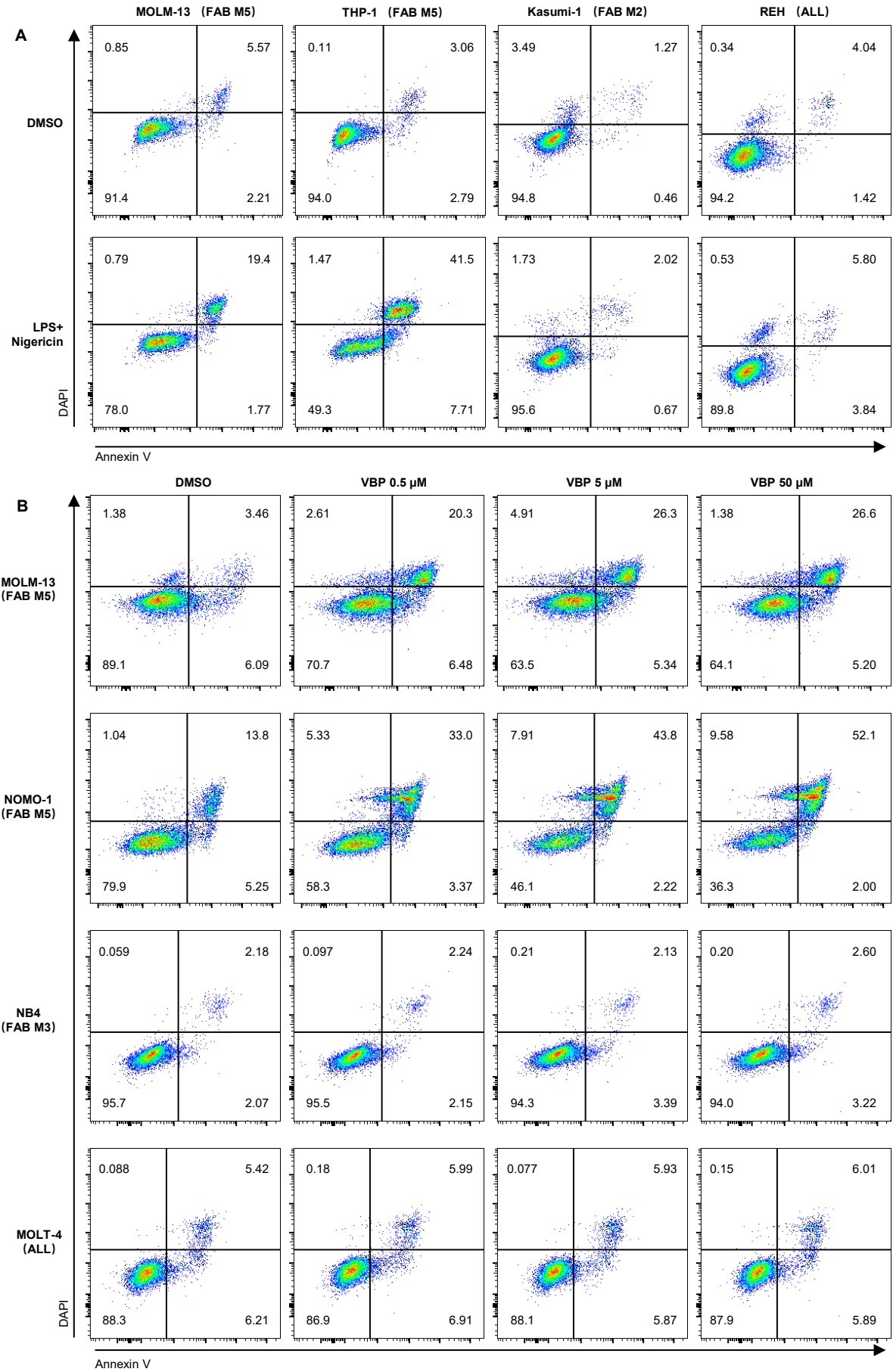

**Supplemental Figure 4. Representative flow-cytometry profiles of leukemia cell death induced by LPS plus nigericin or VBP.**

**(A)** Representative Annexin V/DAPI flow-cytometry plots of the indicated leukemia cell lines—MOLM-13 (FAB M5), THP-1 (FAB M5), Kasumi-1 (FAB M2) and REH (ALL)—following DMSO treatment or LPS priming (10 µg/mL, 24 h) and nigericin treatment (10 µM, 1 h).

**(B)** Representative Annexin V/DAPI flow-cytometry plots of the indicated leukemia cell lines—MOLM-13 (FAB M5), NOMO-1 (FAB M5), NB4 (FAB M3) and MOLT-4 (ALL)—treated with DMSO or VBP (0.5, 5 or 50 µM) for 48 h.

Numbers indicate the percentage of cells in each quadrant. Flow-cytometric quantification of cell death induced by LPS plus nigericin and VBP is shown in Figure 1K and Figure 2C, respectively.

### Supplemental Figure 5

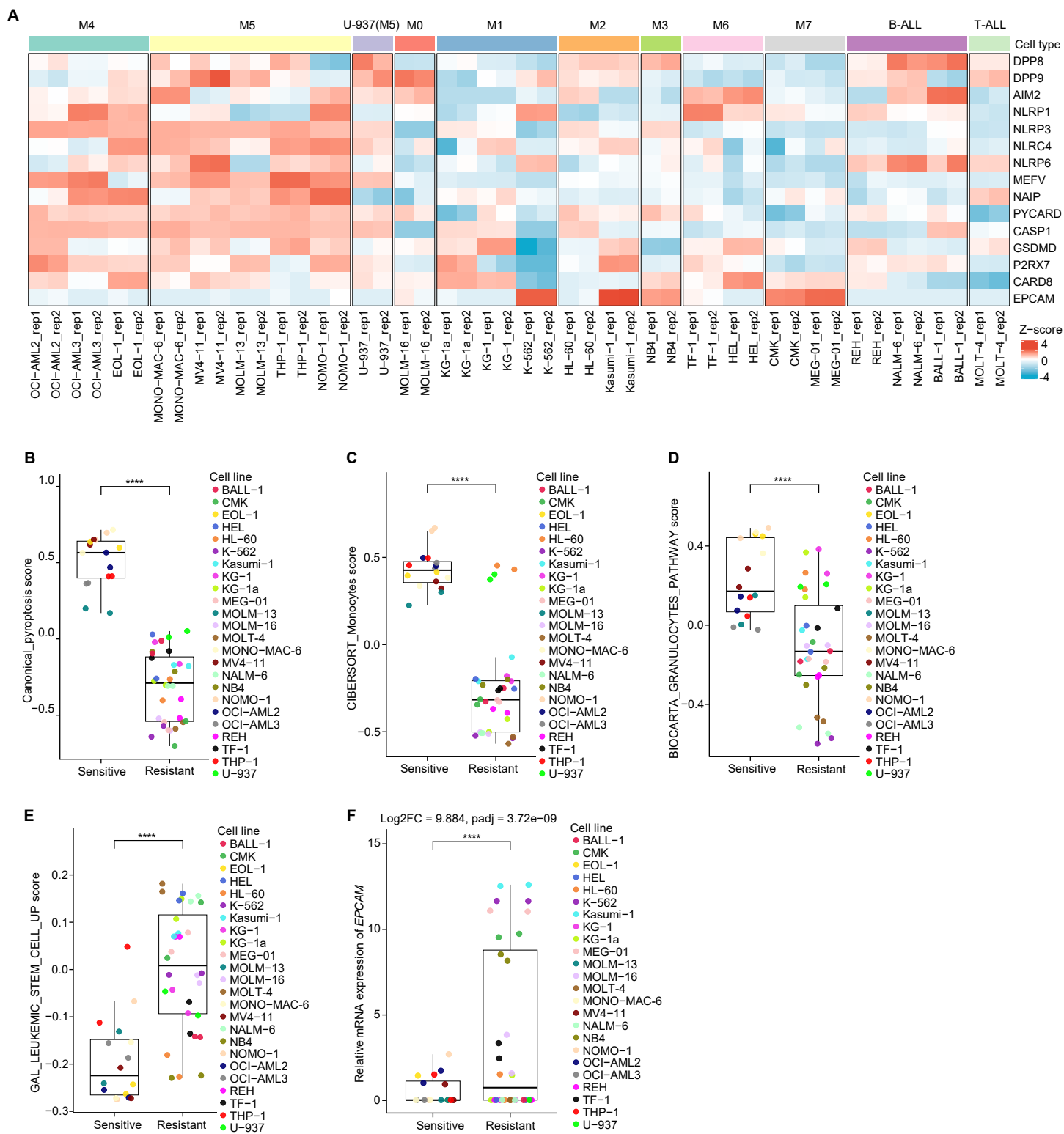

**Supplemental Figure 5. Transcriptomic features associated with VBP sensitivity in leukemia cell lines.**

**(A)** Heat map showing RNA-seq expression of canonical pyroptosis-related genes, together with *EPCAM*, across AML and ALL cell lines. Each cell line is represented by two RNA-seq replicates. Cell lines are ordered by FAB subtype or lymphoid lineage, with U-937 annotated separately. Colors represent row-wise Z-score-normalized expression values.

**(B)** Canonical pyroptosis signature scores in VBP-sensitive and VBP-resistant leukemia cell lines.

**(C)** Monocytic differentiation scores estimated using the CIBERSORT\_Monocytes signature in VBP-sensitive and VBP-resistant leukemia cell lines.

**(D)** Granulocytic differentiation scores based on the BIOCARTE\_GNANULOCYTES\_PATHWAY signature in VBP-sensitive and VBP-resistant leukemia cell lines.

**(E)** Leukemia stem cell signature scores based on the GAL\_LEUKEMIC\_STEM\_CELL\_UP gene set in VBP-sensitive and VBP-resistant leukemia cell lines.

**(F)** Relative *EPCAM* mRNA expression in VBP-sensitive and VBP-resistant leukemia cell lines.

In (B-E), each dot represents one RNA-seq replicate, with two replicates per cell line; boxes indicate the median and interquartile range; whiskers indicate the minimum and maximum values. Statistical significance in (B-E) was determined by two-sided Wilcoxon rank-sum test. In (F), each dot represents one RNA-seq replicate, with two replicates per cell line. Differential expression of *EPCAM* was assessed using DESeq2; log<sub>2</sub> fold change (FC) and adjusted *P* value are indicated. \*\*\**P* < .001, \*\*\*\**P* < .0001.

#### Supplemental Figure 6

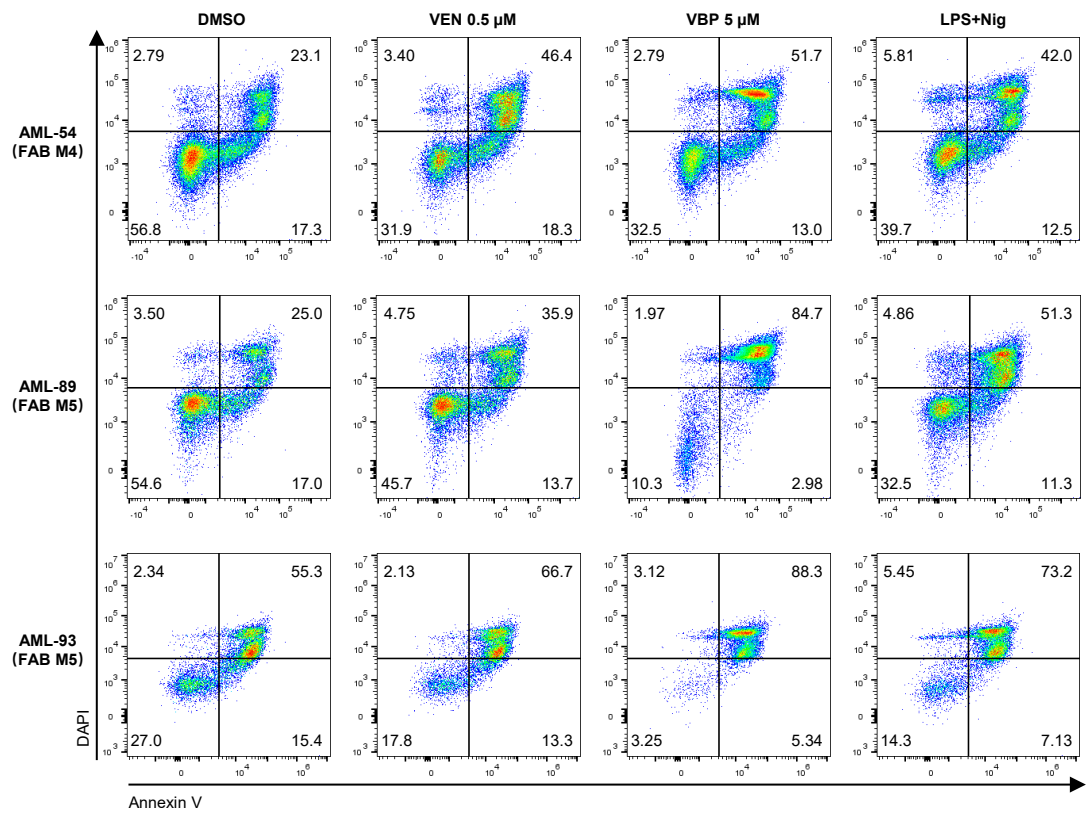

**Supplemental Figure 6. Representative flow-cytometry profiles of primary AML specimens following ex vivo drug treatments.**

Representative Annexin V/DAPI flow-cytometry plots of primary AML BMMNCs from three specimens—AML-54 (FAB M4), AML-89 (FAB M5) and AML-93 (FAB M5)—treated *ex vivo* with DMSO, VEN (0.5 μM, 36 h), VBP (5 μM, 24 h) or LPS+Nig (LPS priming at 10 μg/mL for 24 h followed by nigericin treatment at 10 μM for 1 h). Numbers indicate the percentage of cells in each quadrant. Viable cells were defined as Annexin V<sup>-</sup>DAPI<sup>-</sup>. Flow-cytometry-derived relative viability values following VBP and LPS+Nig treatments are quantified in Figure 3C and Figure 3F, respectively.

### Supplemental Figure 7

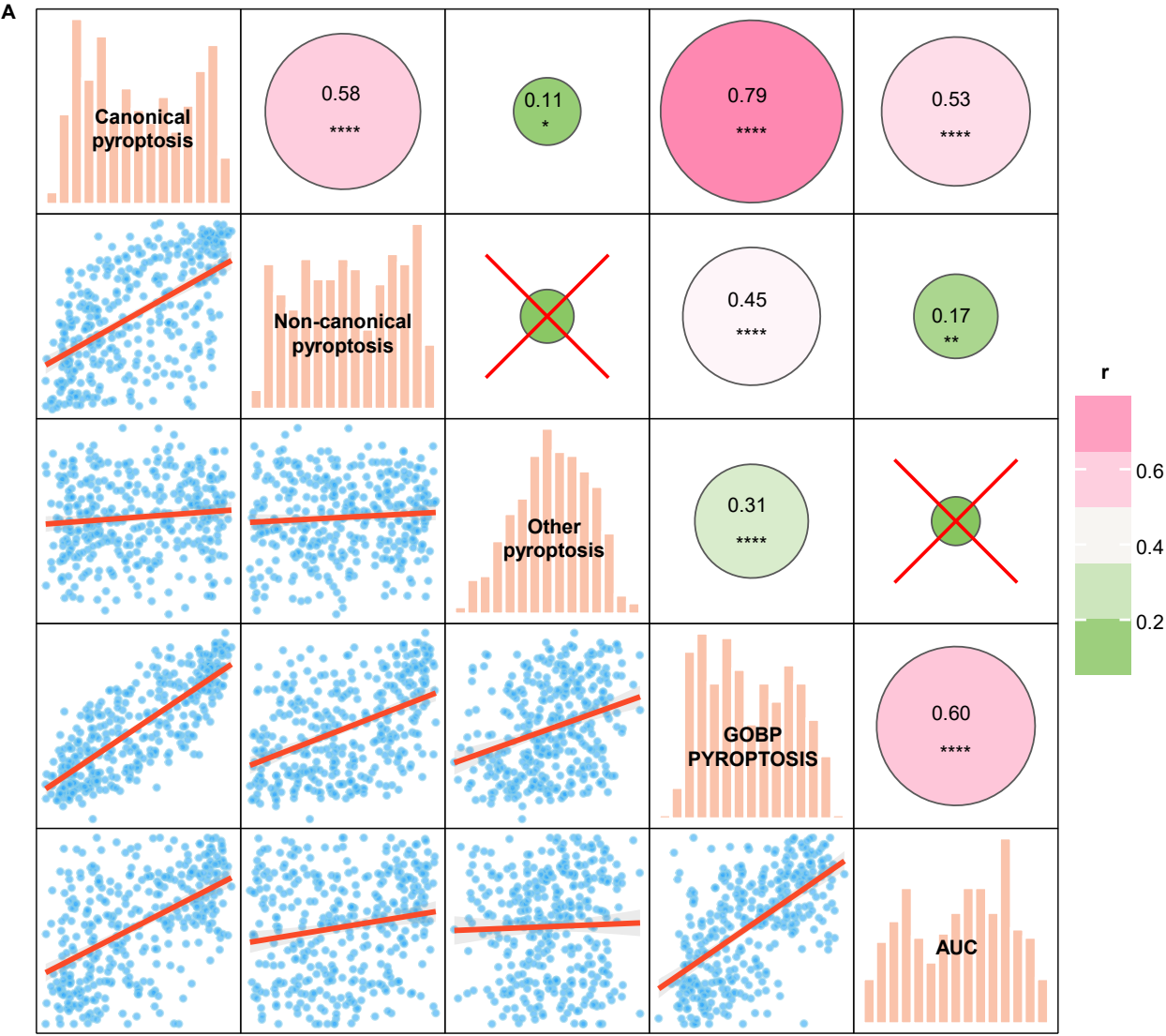

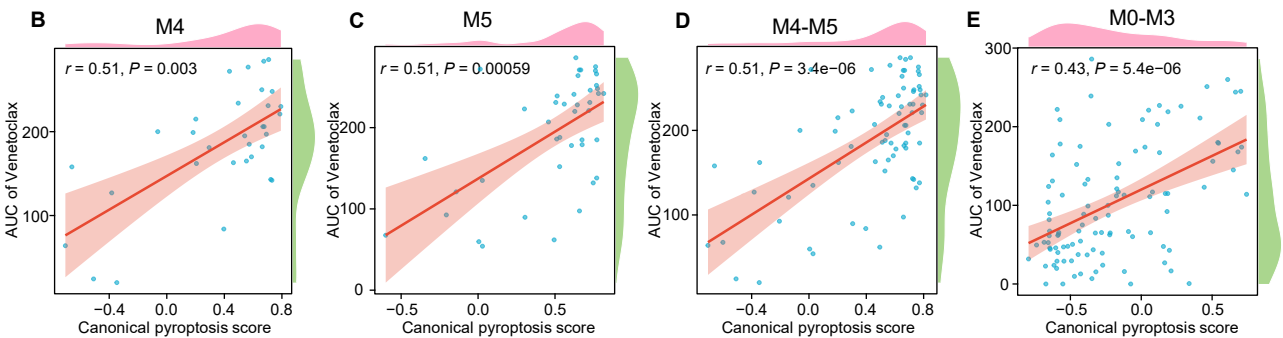

**Supplemental Figure 7. Canonical pyroptosis signature score is positively correlated with VEN resistance in the BeatAML cohort.**

**(A)** Pairwise Spearman correlation matrix of pyroptosis-related signature and gene-set scores and VEN AUC in the BeatAML cohort. Diagonal panels show the distribution of each variable. Lower triangle panels show pairwise scatter plots with fitted regression lines. Upper triangle panels show Spearman correlation coefficients; circle size indicates correlation strength, and circle color indicates corresponding $r$ values according to the scale bar. Red crosses (×) indicate non-significant correlations. \*$P < .05$, \*\*$P < .01$, \*\*\*\*$P < .0001$.

**(B-E)** Spearman correlation between canonical pyroptosis signature score and VEN AUC in AML samples stratified by FAB subtype: M4 (B), M5 (C), combined M4/M5 (D) and M0-M3 (E). Each dot represents one AML sample; fitted lines are shown with 95% confidence intervals. Spearman correlation coefficients and $P$ values are indicated in the plots.

Supplemental Figure 8

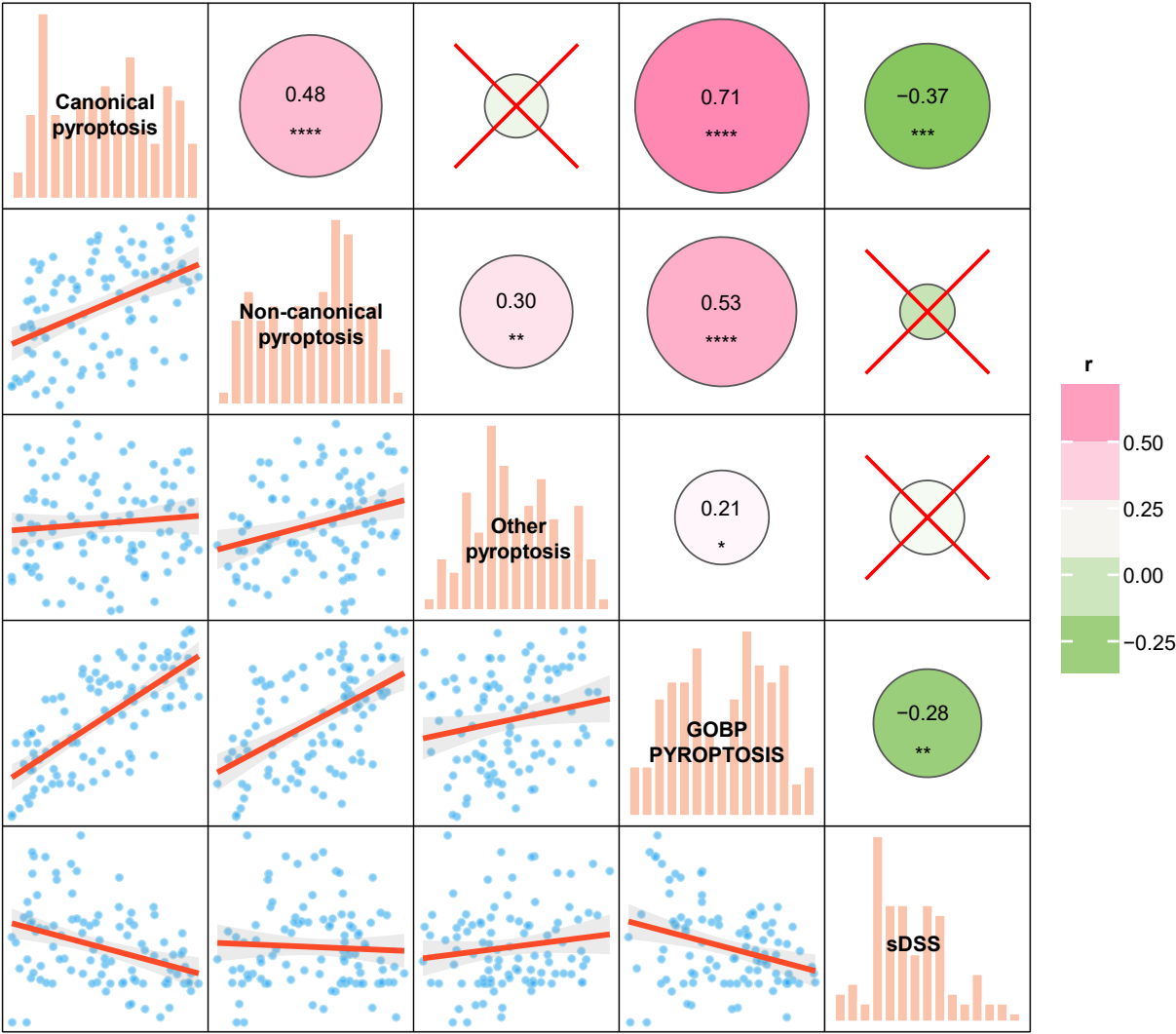

**Supplemental Figure 8. Canonical pyroptosis signature score is negatively correlated with VEN sensitivity in the FIMM-AML cohort.**

Pairwise Spearman correlation matrix of pyroptosis-related signature and gene-set scores and VEN selective drug sensitivity score (sDSS) in the FIMM-AML cohort. Diagonal panels show the distribution of each variable. Lower triangle panels show pairwise scatter plots with fitted regression lines. Upper triangle panels show Spearman correlation coefficients; circle size indicates correlation strength, and circle color indicates corresponding $r$ values according to the scale bar. Each dot represents one AML sample. Red crosses (×) indicate non-significant correlations. $*P < .05$, $**P < .01$, $***P < .001$, $****P < .0001$.

### Supplemental Figure 9

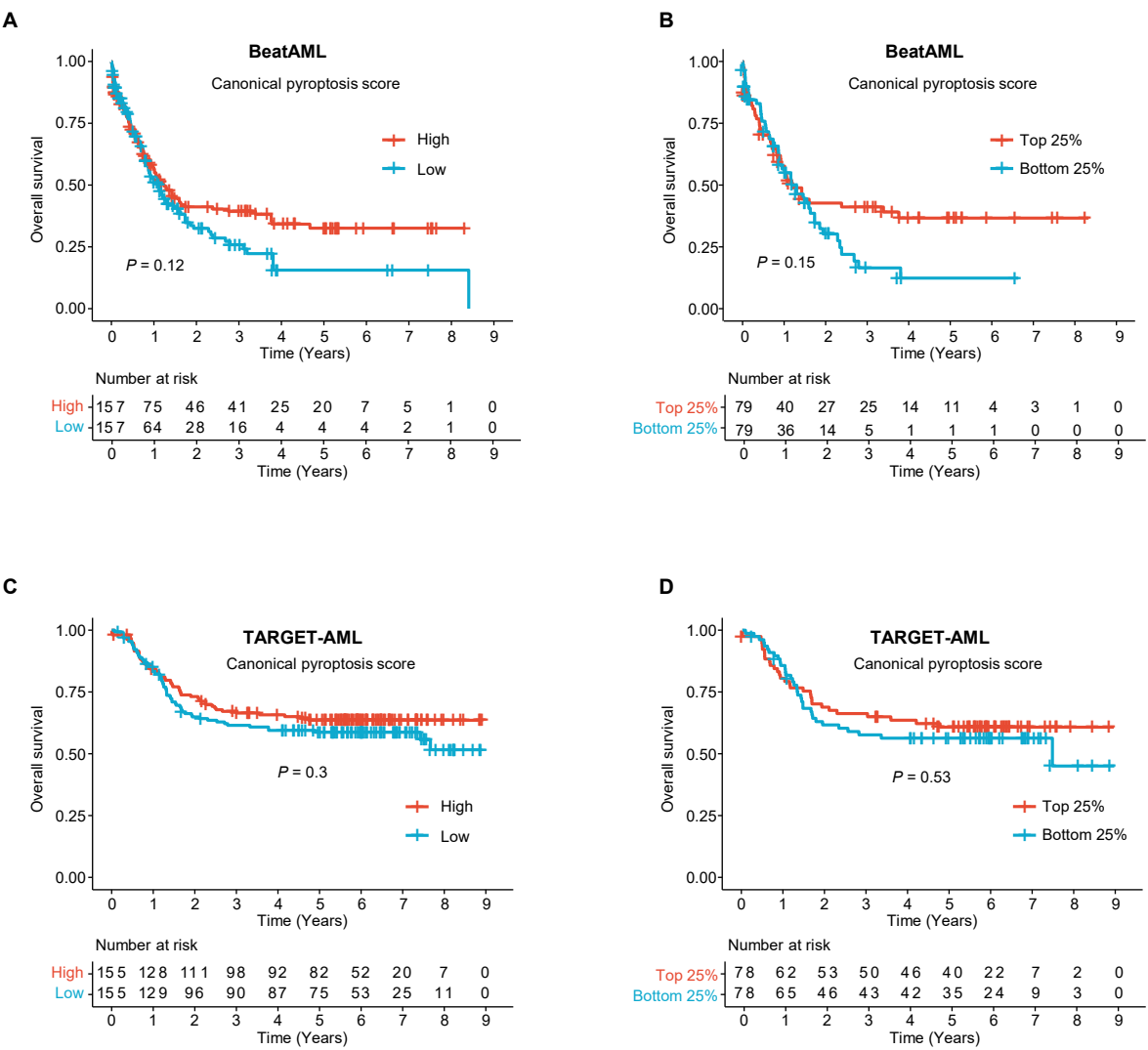

**Supplemental Figure 9. Canonical pyroptosis signature scores are not associated with overall survival in chemotherapy-treated AML cohorts.**

**(A-D)** Kaplan-Meier overall survival curves of chemotherapy-treated AML patients from the BeatAML (A-B) and TARGET-AML (C-D) cohorts, stratified by canonical pyroptosis signature scores. Patients were stratified into high- and low-score groups using the median as the cutoff (A, C), or into top 25% and bottom 25% groups according to the score distribution (B, D). The number of patients at risk is shown below each plot. *P* values were calculated using the log-rank test and are indicated in the plots.

### Supplemental Figure 10

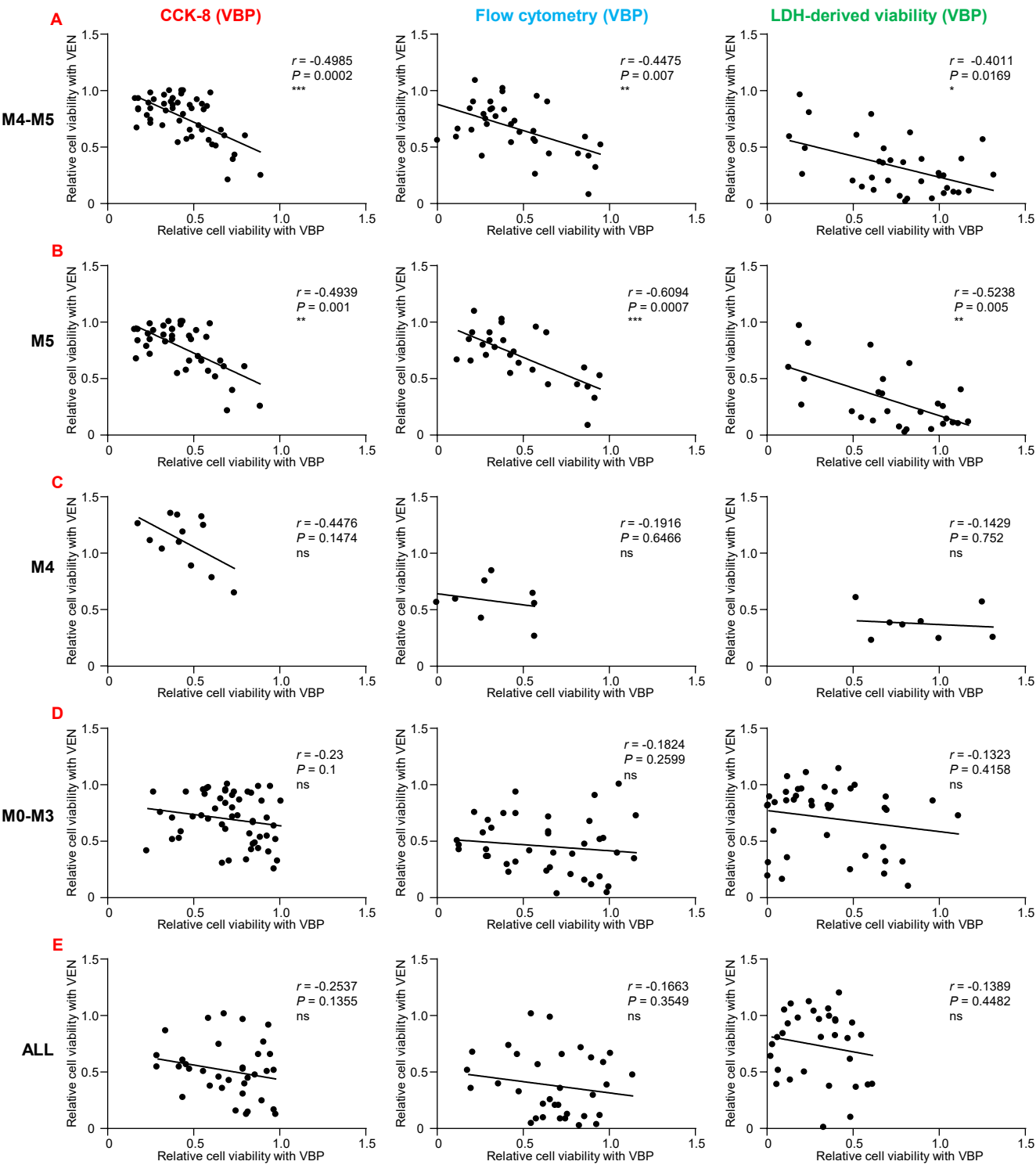

**Supplemental Figure 10. Subtype-specific correlations between VBP and VEN responses in primary leukemia specimens.**

(A-E) Spearman correlations between *ex vivo* responses to VBP (5 μM, 24 h) and VEN (0.5 μM, 36 h) in primary leukemia specimens, assessed by CCK-8 relative viability, relative viability by Annexin V/DAPI flow cytometry and LDH-derived viability. Specimens were stratified as combined FAB M4/M5 AML (A), FAB M5 AML (B), FAB M4 AML (C), FAB M0-M3 AML (D) and ALL (E).

Each dot represents one primary leukemia specimen. Fitted lines are shown, and Spearman correlation coefficients and *P* values are indicated in the plots. Relative viability and LDH-derived viability were calculated as described for Figure 4D-I. ns, not significant; \**P* < .05, \*\**P* < .01, \*\*\**P* < .001, \*\*\*\**P* < .0001.

### Supplemental Figure 11

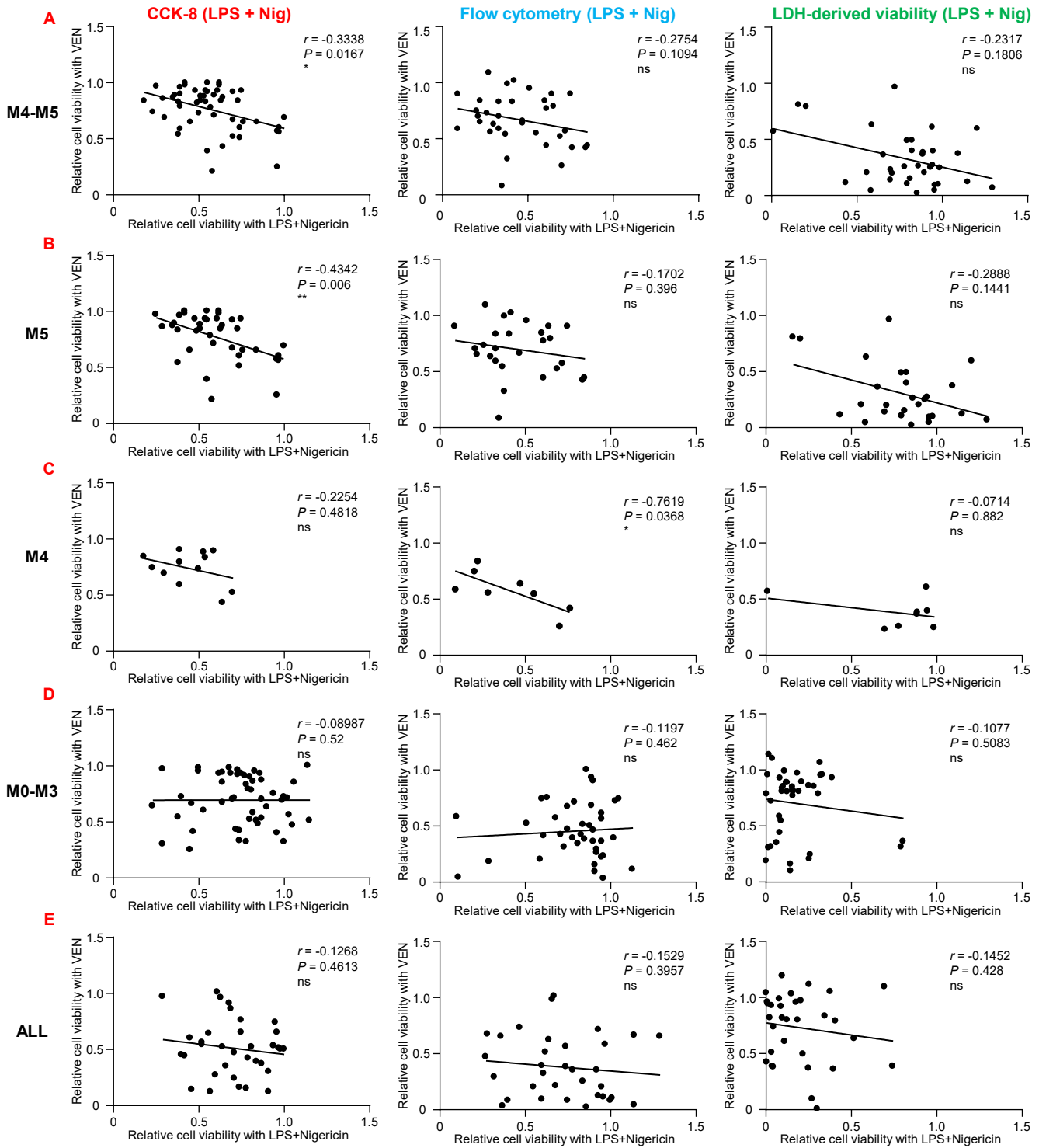

**Supplemental Figure 11. Subtype-specific correlations between LPS+Nig and VEN responses in primary leukemia specimens.**

(A-E) Spearman correlations between *ex vivo* responses to LPS+Nig (LPS, 10 µg/mL, 24 h; nigericin, 10 µM, 1 h) and VEN (0.5 µM, 36 h) in primary leukemia specimens, assessed by CCK-8 relative viability, relative viability by Annexin V/DAPI flow cytometry and LDH-derived viability. Specimens were stratified as combined FAB M4/M5 AML (A), FAB M5 AML (B), FAB M4 AML (C), FAB M0-M3 AML (D) and ALL (E).

Each dot represents one primary leukemia specimen. Fitted lines are shown, and Spearman correlation coefficients and *P* values are indicated in the plots. Relative viability and LDH-derived viability were calculated as described for Figure 4D-I. ns, not significant; \**P* < .05, \*\**P* < .01.

#### Supplemental Figure 12

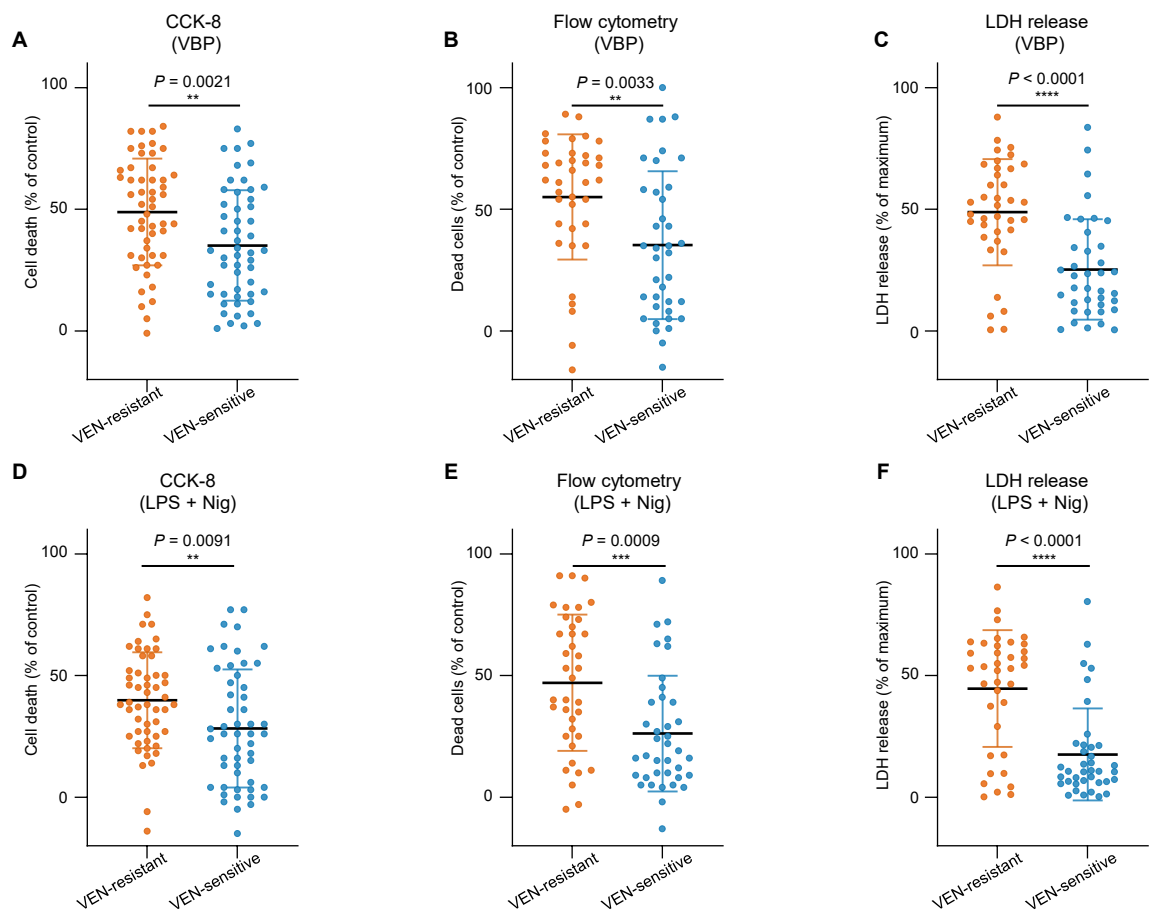

**Supplemental Figure 12. VEN-resistant primary AML specimens show increased sensitivity to canonical pyroptosis induction.**

**(A-C)** *Ex vivo* responses to VBP in VEN-resistant and -sensitive primary AML specimens, assessed by CCK-8-derived cell death (A), Annexin V/DAPI flow-cytometric quantification of dead cells (B) and LDH release (C).

**(D-F)** *Ex vivo* responses to LPS+Nig-induced canonical pyroptosis in VEN-resistant and -sensitive primary AML specimens, assessed by CCK-8-derived cell death (D), Annexin V/DAPI flow-cytometric quantification of dead cells (E) and LDH release (F).

In (A-F), VEN-resistant and VEN-sensitive groups were defined separately for each assay using the median VEN-induced cell death rate among specimens with available data. Drug treatments were performed as described for Figure 3. Each dot represents one primary AML specimen, and data are shown as mean ± s.d. For CCK-8 and flow-cytometry assays, relative viability was normalized to the corresponding DMSO-treated controls, and cell death was calculated as 100% minus relative viability. LDH release is shown as the percentage of maximal release after subtraction of the corresponding DMSO-control baseline. Statistical significance was determined by unpaired two-sided Student's *t*-test. \*\*$P < .01$, \*\*\*$P < .001$, \*\*\*\*$P < .0001$.

### Supplemental Figure 13

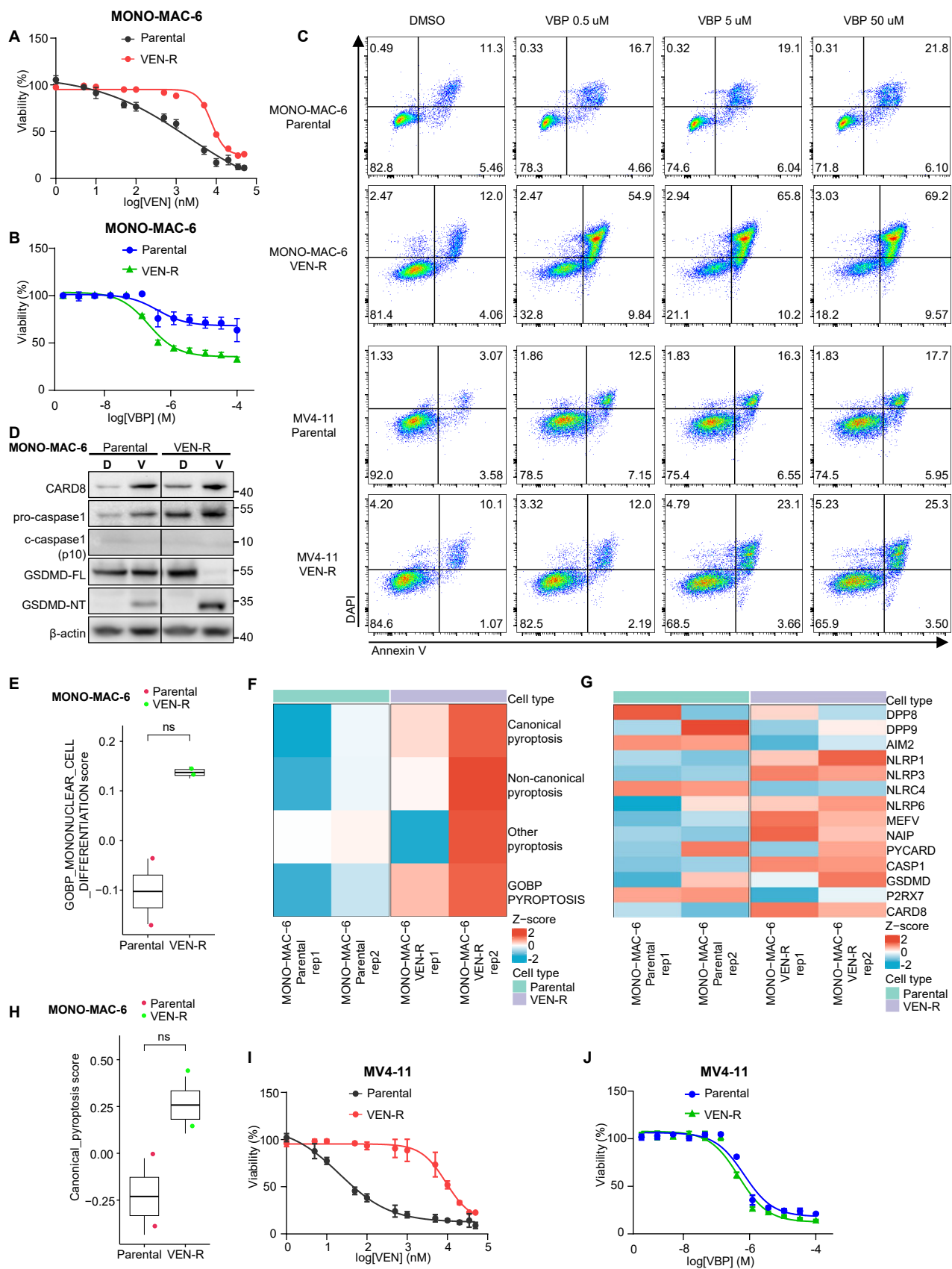

**Supplemental Figure 13. AML cells with acquired VEN resistance retain or exhibit enhanced VBP sensitivity and canonical pyroptotic potential.**

(A) Dose-response curves of VEN in parental and VEN-resistant (VEN-R) MONO-MAC-6 cells.
(B) Dose-response curves of VBP in parental and VEN-R MONO-MAC-6 cells.
(C) Representative Annexin V/DAPI flow-cytometry plots of parental and VEN-R MONO-MAC-6 and MV4-11 cells following treatment with DMSO or the indicated concentrations of VBP for 48 h.
(D) Immunoblot analysis of CARD8, pro-caspase-1, cleaved caspase-1, full-length GSDMD (GSDMD-FL), and the cleaved GSDMD N-terminal fragment (GSDMD-N) in parental and VEN-R MONO-MAC-6 cells following treatment with DMSO (D) or VBP (V; 10 μM, 48 h). β-actin was used as a loading control.
(E) GOBP\_MONONUCLEAR\_CELL\_DIFFERENTIATION scores in parental and VEN-R MONO-MAC-6 cells.
(F) Heat map showing pyroptosis-related signature and gene-set scores in parental and VEN-R MONO-MAC-6 cells, including canonical pyroptosis, non-canonical pyroptosis and other-pyroptosis signatures, together with the GOBP\_PYROPTOSIS gene set.
(G) Heat map showing expression of selected canonical pyroptosis machinery genes in parental and VEN-R MONO-MAC-6 cells.
(H) Canonical pyroptosis signature scores in parental and VEN-R MONO-MAC-6 cells.
(I) Dose-response curves of VEN in parental and VEN-R MV4-11 cells.
(J) Dose-response curves of VBP in parental and VEN-R MV4-11 cells.
- In (A, B, I) and (J), cell viability was normalized to matched DMSO-treated controls, and data are shown as mean ± s.d. from three independent experiments. In (F), colors represent row-wise Z-score-normalized signature or gene-set scores; in (G), colors represent row-wise Z-score-normalized expression values. In (E) and (H), each dot represents one biological replicate; statistical significance was determined by Wilcoxon rank-sum tests. ns, not significant.

Supplemental Figure 14

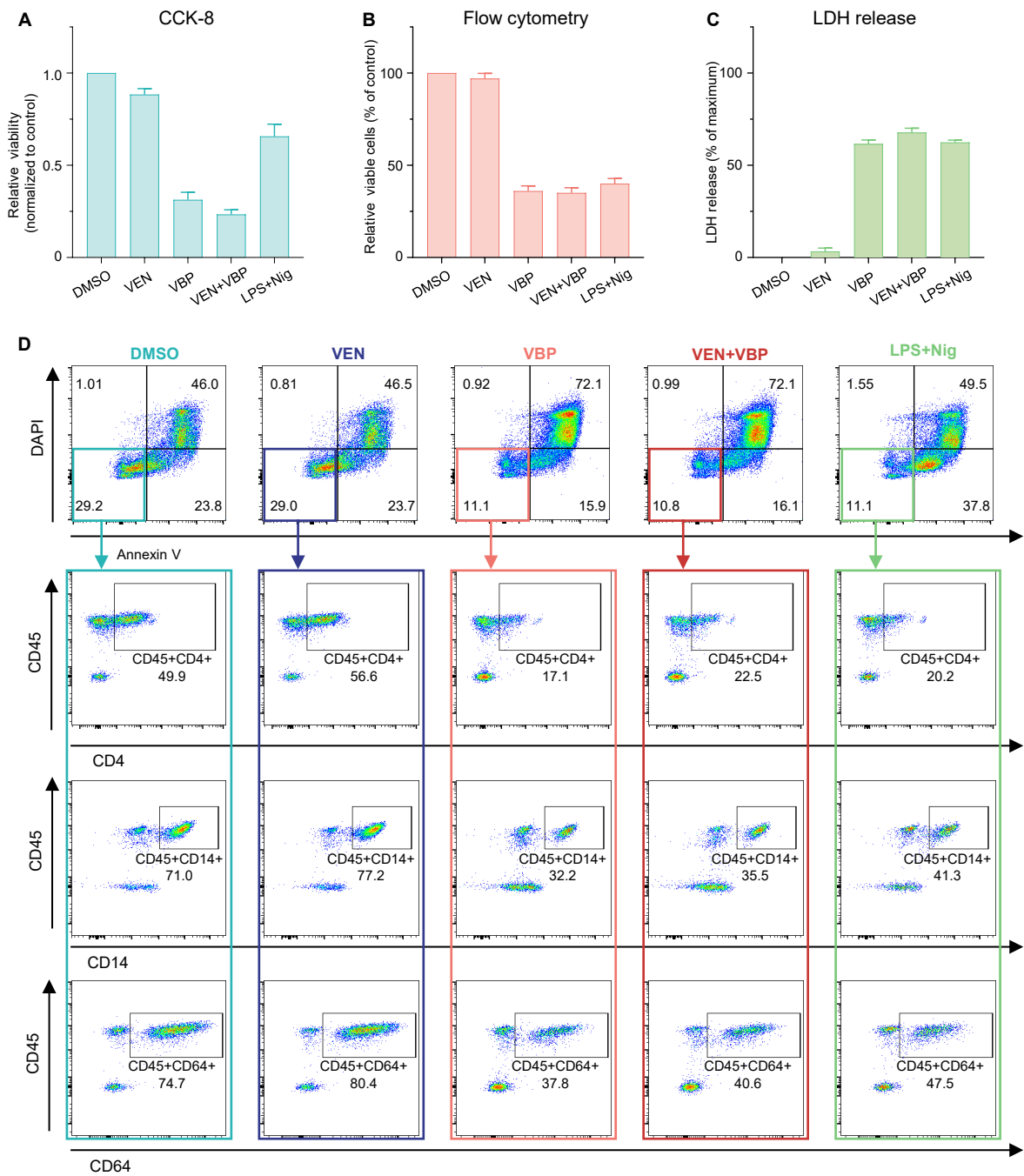

**Supplemental Figure 14. Primary AML-82 cells are resistant to VEN and sensitive to canonical pyroptosis induction *ex vivo*.**

**(A-C)** *Ex vivo* responses of BMMNCs from patient AML-82 treated with DMSO, VEN, VBP, VEN+VBP or LPS+Nig, assessed by CCK-8 relative viability (A), relative viability by Annexin V/DAPI flow cytometry (B) and baseline-corrected LDH release (C).

**(D)** Representative flow-cytometry plots of BMMNCs from patient AML-82 (FAB M5) treated *ex vivo* with DMSO, VEN, VBP, VEN+VBP or LPS+Nig. The top row shows Annexin V/DAPI staining, and the lower three rows show CD45<sup>+</sup>CD4<sup>+</sup>, CD45<sup>+</sup>CD14<sup>+</sup>, and CD45<sup>+</sup>CD64<sup>+</sup> monocytic fractions among gated viable cells defined as Annexin V<sup>-</sup>DAPI<sup>-</sup>. Numbers indicate the percentage of cells in the indicated quadrants or gates.

Drug treatments were performed as described for Figure 3; in the combination group, cells were exposed to VEN for 36 h, with VBP added during the final 24 h. Data in (A-C) are shown as mean ± s.d. of three technical replicates. For (A) and (B), viability was normalized to corresponding DMSO-treated controls. For (C), LDH release was calculated as the percentage of maximal release and corrected by subtraction of the corresponding DMSO-control baseline.

Supplemental Figure 15

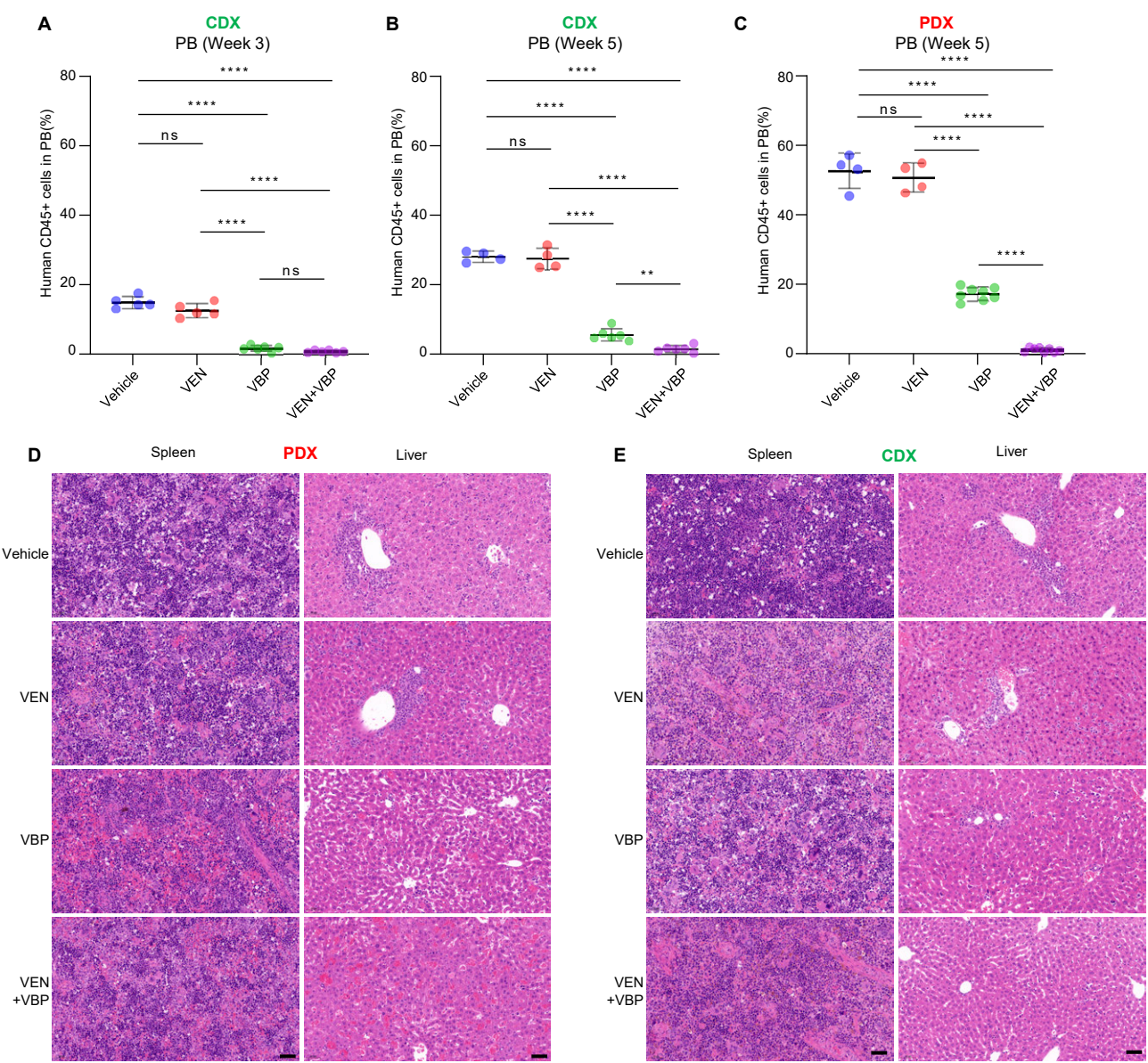

**Supplemental Figure 15. VBP suppresses VEN-resistant monocytic AML progression *in vivo*.**

**(A-C)** Flow-cytometric quantification of hCD45<sup>+</sup> leukemic cells in PB from CDX mice after 3 weeks of treatment (A) and at week 5 (B), and in PB from PDX mice at week 5 (C).

**(D-E)** Representative H&E staining of spleen and liver sections from PDX (D) and CDX (E) mice at week 5 in the indicated treatment groups. Scale bars, 50 μm.

In (A-C), each dot represents one mouse, and data are shown as mean ± s.d. CDX groups initially included $n = 6$ mice each, whereas PDX groups initially included $n = 8$ mice each; at week 5, group sizes varied because mice reaching humane endpoints before the scheduled terminal collection were unavailable for terminal analyses.

Statistical significance was determined by one-way ANOVA followed by Tukey's multiple-comparisons test for indicated comparisons. ns, not significant; $*P < .05$, $**P < .01$, $****P < .0001$.

#### **Descriptions for Separate Supplemental Tables**

##### **Supplemental Table 1. Publicly available datasets analyzed in this study.**

Dataset accession identifiers, experimental platforms, cohort descriptions, source publications, journals, and DOIs are provided for the microarray, bulk RNA-seq, and single-cell RNA-seq datasets used in this study.

##### **Supplemental Table 2. Pyroptosis-related gene signatures and gene sets used in this study.**

The biological definition, source, version, number of genes, and gene symbols are provided for the canonical pyroptosis, non-canonical pyroptosis, other-pyroptosis, and GOBP\_PYROPTOSIS gene sets used in the analyses.

##### **Supplemental Table 3. Clinical and molecular characteristics of primary leukemia specimens analyzed in this study.**

The Cohort summary worksheet summarizes demographic, diagnostic, hematologic, cytogenetic, and recurrent molecular characteristics of the 142 primary leukemia specimens (106 AML and 36 ALL). The Sample metadata worksheet provides individual-level clinical, cytogenetic, mutation, and fusion annotations for the analyzed specimens.
